# Gene silencing by double-stranded RNA from *C. elegans* neurons reveals functional mosaicism of RNA interference

**DOI:** 10.1101/393074

**Authors:** Snusha Ravikumar, Sindhuja Devanapally, Antony M Jose

**Affiliations:** Department of Cell Biology and Molecular Genetics, University of Maryland, College Park, MD-20742.

**Keywords:** RNA-dependent RNA polymerase, stochastic, systemic RNAi

## Abstract

Delivery of double-stranded RNA (dsRNA) into animals can silence genes of matching sequence in diverse cell types through mechanisms that have been collectively called RNA interference. In the nematode *C. elegans*, dsRNA from multiple sources can trigger the amplification of silencing signals. Amplification occurs through the production of small RNAs by two RNA-dependent RNA polymerases (RdRPs) that are thought to be tissue-specific - EGO-1 in the germline and RRF-1 in somatic cells. Here we analyze instances of silencing in somatic cells that lack RRF-1. By varying dsRNA sources and target genes, we find that silencing in the absence of RRF-1 is most obvious when dsRNA from neurons is used to silence genes in intestinal cells. This silencing requires EGO-1, but the lineal identity of cells that can use EGO-1 varies. This variability could be because random sets of cells can either receive different amounts of dsRNA from the same source or use different RdRPs to perform the same function. As a result, all cells appear similarly functional despite underlying differences that vary from animal to animal. This functional mosaicism cautions against the use of a few molecules as proxies for predicting the behavior of a cell.

**Graphical Abstract:** 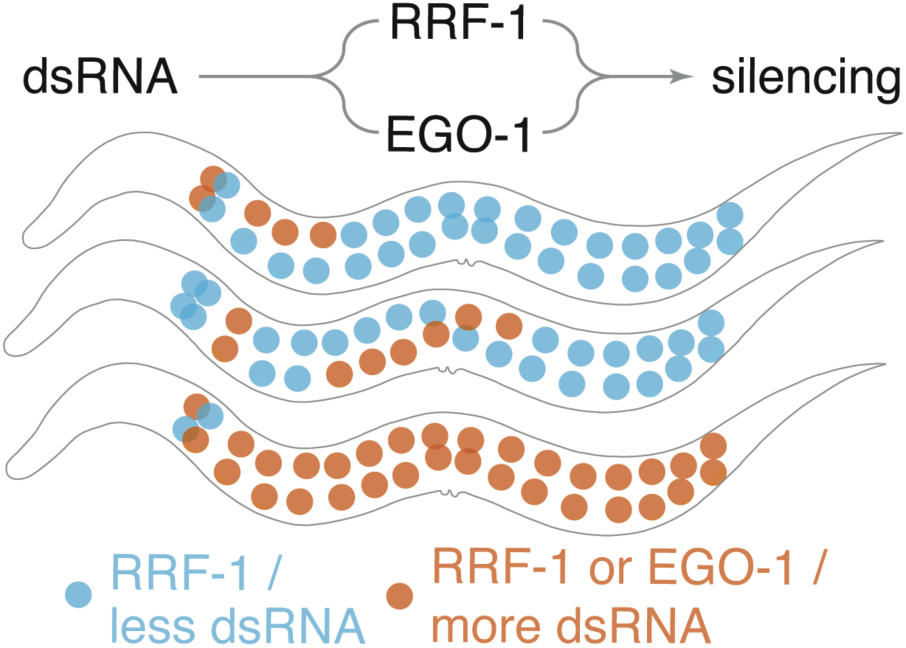

Random sets of cells can either receive different amounts of double-stranded RNA from neurons or use different RdRPs – RRF-1 only versus RRF-1 or EGO-1 – to perform the same function.

## INTRODUCTION

Animals have diverse cell types that perform specialized functions while retaining the ability to perform common functions. Such common functions could rely on the same molecular machinery in all cells or on different machinery in different cells. As a result, an apparently uniform organismal response could obscure differences in the mechanisms used by different cells. A common response to viral infection is the silencing of viral genes facilitated by the recognition of viral double-stranded RNA (dsRNA) (reviewed in (1)). The experimental addition of dsRNA triggers similar mechanism(s) that can silence any matching sequence (2). This process of RNA interference is a powerful approach for gene silencing applications in a variety of organisms (reviewed in (3)). In the nematode *C. elegans*, exposure to different sources of dsRNA can silence matching genes in many somatic cell types and in the germline (4–6). Studies in *C. elegans* have therefore been informative in piecing together the organismal response to RNAi in an animal. While similar silencing responses occur in diverse cell types, it is unclear whether dsRNA from every source engages the same molecular machinery in each cell.

Entry of extracellular dsRNA into the cytosol and subsequent silencing relies on the conserved dsRNA importer SID-1 (7–10). SID-1-dependent silencing is observed in many tissues even when dsRNA is expressed within a single tissue, suggesting that form(s) of dsRNA move between cells. In particular, dsRNA expressed in neurons can silence a target gene in somatic tissues such as the intestine, muscle, and hypodermis (11–13) and in the germline (14). Silencing in these diverse target cells requires the dsRNA-binding protein RDE-4 (15, 16) and the endonuclease DCR-1, which together process dsRNA into small-interfering RNAs (siRNAs) (17, 18), and the Argonaute RDE-1, which binds siRNAs (19). Upon recognition of a matching mRNA by RDE-1-bound siRNAs, RNA-dependent RNA Polymerases (RdRPs) are recruited, resulting in the production of numerous secondary siRNAs (20, 21). Testing multiple target genes suggests that two different RdRPs are used for silencing: RRF-1 for genes expressed in somatic cells (20–22) and EGO-1 for genes expressed in the germline (23, 24). Secondary siRNAs can bind the Argonaute NRDE-3 in somatic cells (25) or the Argonaute HRDE-1 in the germline (26–28) and subsequently accumulate within the nuclei of cells that express the target gene. Through these events, extracellular dsRNA can reduce the levels of mRNA and/or pre-mRNA of a target gene.

While silencing by all extracellular dsRNA requires SID-1, DCR-1, and RDE-1, the requirement for other components can vary. For example, some genes expressed in somatic cells can be silenced by ingested dsRNA in the absence of RRF-1 (29). While many genes do not require NRDE-3 for silencing, the *bli-1* gene requires NRDE-3 for silencing by ingested dsRNA or neuronal dsRNA (13). Finally, a strict requirement for NRDE-3 but not for RRF-1 is seen for the silencing of repetitive DNA that occurs in an enhanced RNAi background upon growth at lower temperatures (30). These observations suggest that a mix of mechanisms could underlie RNAi in *C. elegans.* Experiments that control one variable at a time are needed to elucidate features that dictate the choice of mechanism used for silencing.

Here we reveal that silencing by neuronal dsRNA can differ from silencing by other sources of dsRNA in its requirement for EGO-1 in the absence of RRF-1. We provide a single-cell resolution view of silencing by neuronal dsRNA and find that each animal has a different set of intestinal cells that can rely on EGO-1 for gene silencing.

## MATERIALS AND METHODS

### Strains and Oligonucleotides Used

All strains (listed in Supplementary Table S1) were cultured on Nematode Growth Medium (NGM) plates seeded with 100 μl of OP50 at 20°C and mutant combinations were generated using standard methods (31). Reference alleles indicated as *gene(-)* are as follows: *eri-1(mg366), rrf-1(ok589)*, *rde-1(ne219)*, *rde-11(hj37)*, *sid-1(qt9)*, and *mut-16(pk710)*. Sequences of oligonucleotides used to genotype different mutant combinations are in Supplementary Table S2 (*eri-1*: P01-P02, *rde-1*: P03-P04, *rde-11*: P05-P06, *sid-1*: P07-P08, *rrf-1*: P09-P11, *mut-2/rde-3*: P12-P13, and *mut-16*: P14-P15).

### Transgenesis

*C. elegans* was transformed with plasmids and/or PCR products using microinjection (32) to generate extrachromosomal or integrated arrays. pHC337 was used to express an inverted repeat of *gfp* in neurons (11), which is expected to generate a hairpin RNA (*gfp-*dsRNA). Generation of the array that expresses *unc-22*-dsRNA in neurons (*qtEx136)* was described earlier (12). To rescue silencing defects in *rde-1(jam1)* and *rrf-1(jam3)* animals (Supplementary Figure S2), genomic DNA from wild-type animals (N2 gDNA) was used as a template to generate fused promoter/gene products through overlap extension PCR using Expand Long Template polymerase (Roche) and PCR products were purified using QIAquick PCR Purification Kit (Qiagen). The plasmid pHC448 for *DsRed2* expression in the pharynx or a PCR product, *Prgef-1::DsRed2::unc-54 3’UTR*, for *DsRed2* expression in neurons was used as a co-injection marker (12). Additional details are provided in Supplementary Materials and Methods.

### Genome editing

Synthetic CRISPR RNA (crRNA) and trans-activating crRNA (tracrRNA) (IDT) or single guide RNAs (sgRNA) transcribed in vitro were combined with Cas9 protein (PNA Bio Inc. or IDT) to generate complexes used for genome editing. To transcribe guide RNAs, the scaffold DNA sequence was amplified from pDD162 (*Peft-3::Cas9* + *dpy-10* sgRNA - Addgene plasmid # 47549, a gift from Bob Goldstein) (33) using a common reverse primer (P16) and target-specific forward primers (see Supplementary Table S2), purified (PCR Purification Kit, Qiagen), and used for in vitro transcription (SP6 RNA polymerase, NEB). Deletions were made using two guide RNAs and a single-stranded DNA oligonucleotide repair template with a co-conversion strategy (34). Insertions of *gfp* were performed using a single guide RNA and a double-stranded repair template amplified using PCR (35). *Punc-22::unc-22::gfp* resulted in GFP fluorescence within the pharynx as reported earlier (36). Additional details are provided in Supplementary Materials and methods.

### Feeding RNAi

One generation of feeding RNAi was performed as described earlier (9) and the numbers of brightly fluorescent intestinal nuclei in animals subject to RNAi were counted for Figure 1D.

**Figure 1:**
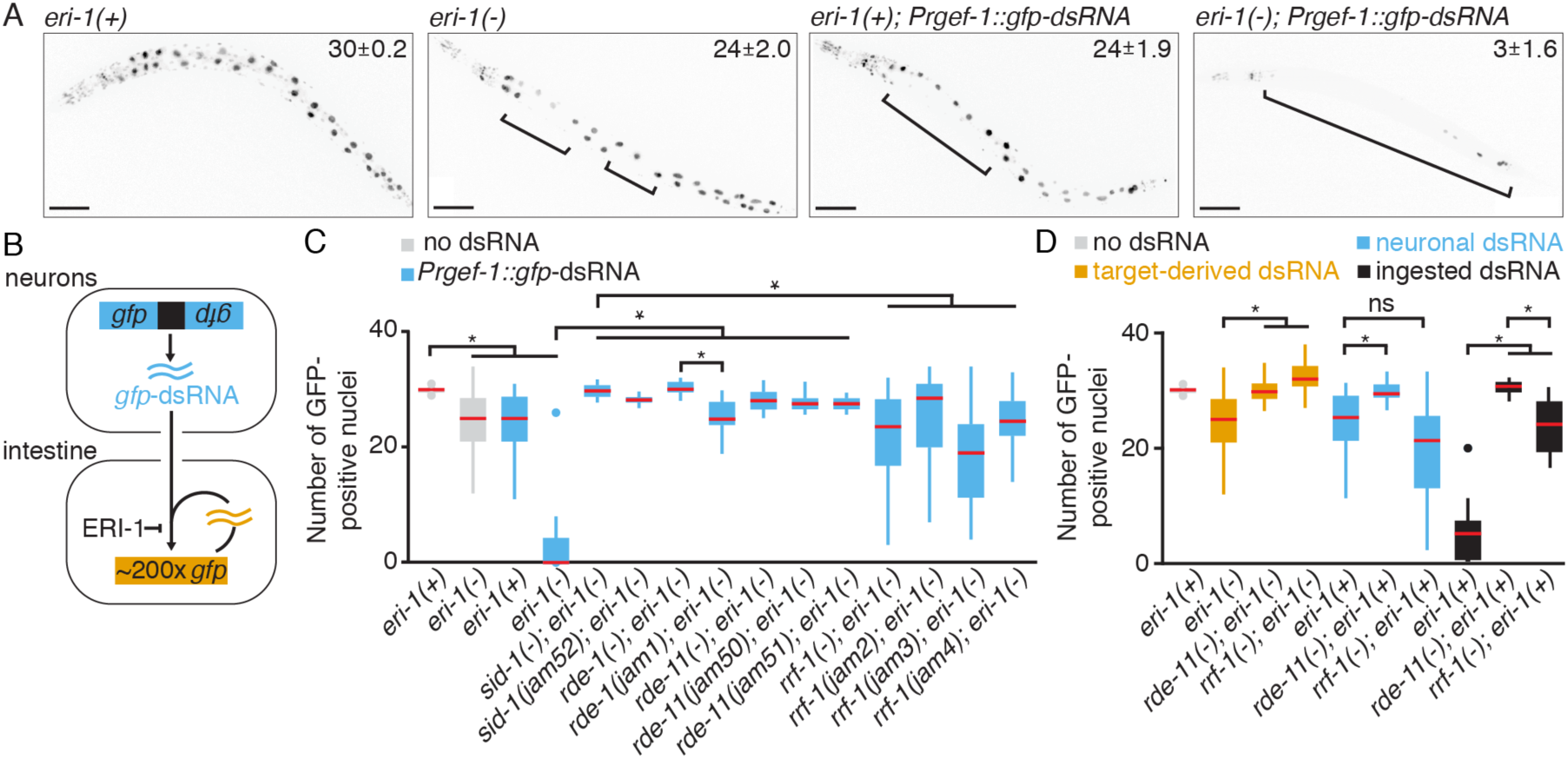
Silencing by different sources of double-stranded RNA show synergy and can have different requirements for the RNA-dependent RNA polymerase RRF-1. (**A**) Silencing upon loss of *eri-1* and by neuronal dsRNA shows synergy. Representative L4-staged animals that express GFP (black) in all tissues (*sur-5::gfp*) in *eri-1(+)* (i.e., wild-type) or *eri-1(-)* backgrounds and animals that in addition express dsRNA against *gfp* in neurons (*Prgef-1::gfp*-dsRNA) in either background are shown. Brackets indicate silenced intestinal nuclei. Average numbers of GFP positive intestinal nuclei are indicated with 95% confidence intervals (n = 20 animals). Scale bar = 50 µm. (**B**) Schematic of *gfp* silencing in intestinal cells. Silencing by neuronal dsRNA (blue) and by dsRNA made from a multicopy *sur-5::gfp* transgene (orange) are both inhibited by the endonuclease ERI-1. (**C**) Combined silencing by the two sources of dsRNA is strictly dependent on *sid-1*, *rde-1*, and *rde-11*, but partially dependent on *rrf-1*. Silencing of *sur-5::gfp* was measured by counting the number of GFP-positive intestinal nuclei in animals expressing no dsRNA in an *eri-1(+)* or *eri-1(-)* background, in animals expressing *Prgef-1::gfp-*dsRNA in an *eri-1(+)* or *eri-1(-)* background, and in animals expressing *Prgef-1::gfp-*dsRNA in an *eri-1(-)* background with additional mutations in *sid-1*, *rde-1*, *rde-11 or rrf-1.* Known null alleles are represented as *gene(-)* (see Materials and Methods for allele names) and alleles isolated in the screen are represented as *gene(jam#)*. Red bars indicate medians, n ≥ 20 L4-staged animals and asterisks indicate p-value <0.05 (Student’s t-test). (**D**) Unlike silencing by target-derived dsRNA or ingested dsRNA, silencing by neuronal dsRNA is partially independent of RRF-1 and strongly dependent on RDE-11. Silencing was separately measured for the three sources of dsRNA: target-derived dsRNA upon loss of *eri-1* in *eri-1(-)*, *eri-1(-); rde-11(-)* or *eri-1(-); rrf-1(-)* animals (orange), neuronal dsRNA upon expression of *Prgef-1::gfp*-dsRNA in *eri-1(+)*, *eri-1(+); rde-11(-)* or *eri-1(+); rrf-1(-)* animals (blue), or ingested dsRNA from bacteria expressing *gfp*-dsRNA in *eri-1(+)*, *eri-1(+); rde-11(-)*, or *eri-1(-) rrf-1(-)* animals (black). Red bars, n, and asterisks are as in C, and ns = not significant.

### Genetic screen and whole genome sequencing

AMJ1 animals were mutagenized with 25 mM N-ethyl N-nitrosourea (ENU, Toronto Research Chemicals) and ∼600,000 of their F2 progeny were screened for recovery of GFP expression in intestinal cells (performed by A.M.J. in Craig Hunter’s lab, Harvard University). For 23 mutants that showed different degrees of fluorescence, we prepared genomic DNA from ∼1-2 ml of worms (200 - 800 ng/µl of DNA per mutant, NanoVue Plus (GE)). Libraries for Illumina sequencing were prepared at the IBBR sequencing core as per manufacturer’s instructions and sequenced using a HiSeq1000 (Illumina).

### Bioinformatic Analysis

All bioinformatic analyses were done using the web-based Galaxy tool collection (https://usegalaxy.org) (37–39). For each of the 23 mutant strains, we obtained ∼40 million 101 base fastq reads on average. One 5’-end base and three 3’- end bases were of lower quality and were trimmed from all reads before alignment to ce6/WS190 using Bowtie (∼36 million mapped reads per mutant on average). Sequence variants were filtered to call mutations (Phred33 ≥ 20, ≥ 2 aligned reads, and same variant call in ≥ 66% of reads). We intended to rescue any *sid-1* mutations that might arise in the screen to avoid isolating many alleles of *sid-1* (∼100 alleles of *sid-1* were isolated in the original *sid* screen). However, our sequencing data reveals that we had instead inadvertently introduced a non-functional copy with 12 missense mutations as part of the *qtIs50* transgene (Supplementary Figure S1C). Therefore, the threshold for calling a mutation was reduced from 66% to 15% for *sid-1* sequences. For all mutants, non-synonymous changes, changes in splice junctions, and deletions (characterized by lower than average coverage) were analyzed further. Identical changes detected in two or more mutants were eliminated as potential background mutations that were likely present before mutagenesis. Pairwise comparisons were carried out between all mutants to identify cases of different mutations in the same gene (i.e. in silico complementation (40)). Because this process entails 253 pairwise comparisons, we expect that one or two such shared genes will be identified for some mutant pairs at random. For example, for mutant pairs with 30 mutated genes each, the p-value for one shared gene (0.044) and that for two shared genes (0.0009) are both larger than the Bonferroni corrected p-value of 0.0002 for 253 comparisons at α = 0.05 (41).

### Single-molecule RNA fluorescence in situ hybridization (smFISH)

smFISH was performed as described earlier (42, 43). Briefly, custom Stellaris probes recognizing exons of *gfp* (probes spanning exon-exon junctions were not included) labeled with Quasar 670 dye (Biosearch Technologies) were added to fixed L4-staged animals. RNA hybridization was performed with 0.025 µM of probe mix for 48 hours at 37°C in 100 µl of hybridization buffer (10% dextran sulphate (w/v), 2x saline-sodium citrate (SSC), 10% formamide (v/v)). Following a wash in wash buffer (2x SSC, 10% formamide, 0.1% Tween-20) samples were stained with DAPI (4′,6-diamidino-2-phenylindole) for 2 hours at room temperature and washed 5 more times. Before imaging, samples were stored in GLOX (2x SSC, 0.4% glucose, 0.01M Tris, pH 8.0) buffer at 4°C for fewer than 3 hours. Samples were mounted in 10 µl of GLOX buffer and enzymes (glucose oxidase, catalase, and 6-hydroxy-2,5,7,8-tetramethylchroman-2-carboxylic acid (trolox)) and coverslips were sealed with a melted mixture of vaseline, lanolin and paraffin.

### Western blotting

Mixed stage animals were washed off three to five 100mm plates and used for western blot analysis. Samples were sonicated four times (40% amplitude with 45 second pauses between 15 second pulses) using a probe sonicator with a microtip (Branson Sonifier). Proteins were separated on a 14% SDS-PAGE and then blotted onto nitrocellulose paper (TransBlot™ Turbo Midi transfer pack). The blot was probed for GFP first, stripped (incubated in 0.2% Sodium dodecyl sulfate, 0.1 M Tris, pH 8.0, and 1.4% β-mercaptoethanol for 1 hr at 65°C), and then probed for Tubulin. The following primary antibodies were used: mouse anti-α-Tubulin (Sigma: T5168; 1:4000 dilution) and mouse anti-GFP (Thermo Fisher Scientific: MA5-15256; 1:2000 dilution). The following corresponding secondary antibodies were used: Rabbit anti-mouse IgG1 HRP (Sigma: SAB3701171, 1:250 dilution) and goat anti-Mouse IgG(H+L) HRP (Thermo Fisher Scientific: 32430, 1:750 dilution). Blots were developed using chemiluminescence detection reagents (Thermo Fisher Scientific: SuperSignal™ West Pico PLUS) and imaged using a ChemiDoc (Bio-Rad). The western blots in Supplementary Figure S3D are representative of three technical replicates. Signal of the band of interest was quantified using FIJI (NIH, (44)) and is reported as median of ratios with respect to α-tubulin.

### Microscopy

For Figures 2A and 3A and Supplementary Figures S3B, S5, S6B, S6D and S7, animals were immobilized in 5µl of 3mM levamisole (Sigma-Aldrich; catalog no. 196142), mounted on slides, and imaged using an AZ100 microscope (Nikon) at a fixed magnification under non-saturating conditions of the tissue being quantified for silencing. A C-HGFI Intensilight Hg Illuminator was used to excite GFP (filter cube: 450-490 nm excitation, 495 nm dichroic, 500-550 nm emission), which also resulted in some bleed through from the DsRed fluorescence (e.g. Figure 3A). For Figure 4A-4D and Supplementary Figure S8, L4-staged worms were mounted onto a slide with a 3.5% agarose pad after incubating the worm for 10 minutes in 7µl of 1mM freshly made levamisole. Extended exposure to levamisole was necessary for reliable immobilization of the worm for the ∼100 minutes of imaging that was required to obtain 512 x 512 images of entire L4-staged *sur-5::gfp* worms using a 63x lens in a Leica SP5X confocal microscope (average of 3 measurements per line, 319 slices per section, 5 sections, and 0.125 µm between slices). A 488 nm laser was used to excite GFP (emission: 498-550nm, NA=1.4). For Figure 4E and 4F, DAPI, GFP and Quasar 670 fluorescence in intestinal cells anterior to the germline and posterior to the pharynx was acquired as 1024 x 1024 images (6 slices, 0.5 µm between slices) using a 63x lens and 2x digital zoom in a Leica SP5X confocal microscope. GFP was excited as described above, a 405 diode laser was used to excite DAPI (emission: 422-481nm, 9% power) and a 633nm laser was used to excite Quasar 670 (emission: 650-715nm, 50% power).

**Figure 2:**
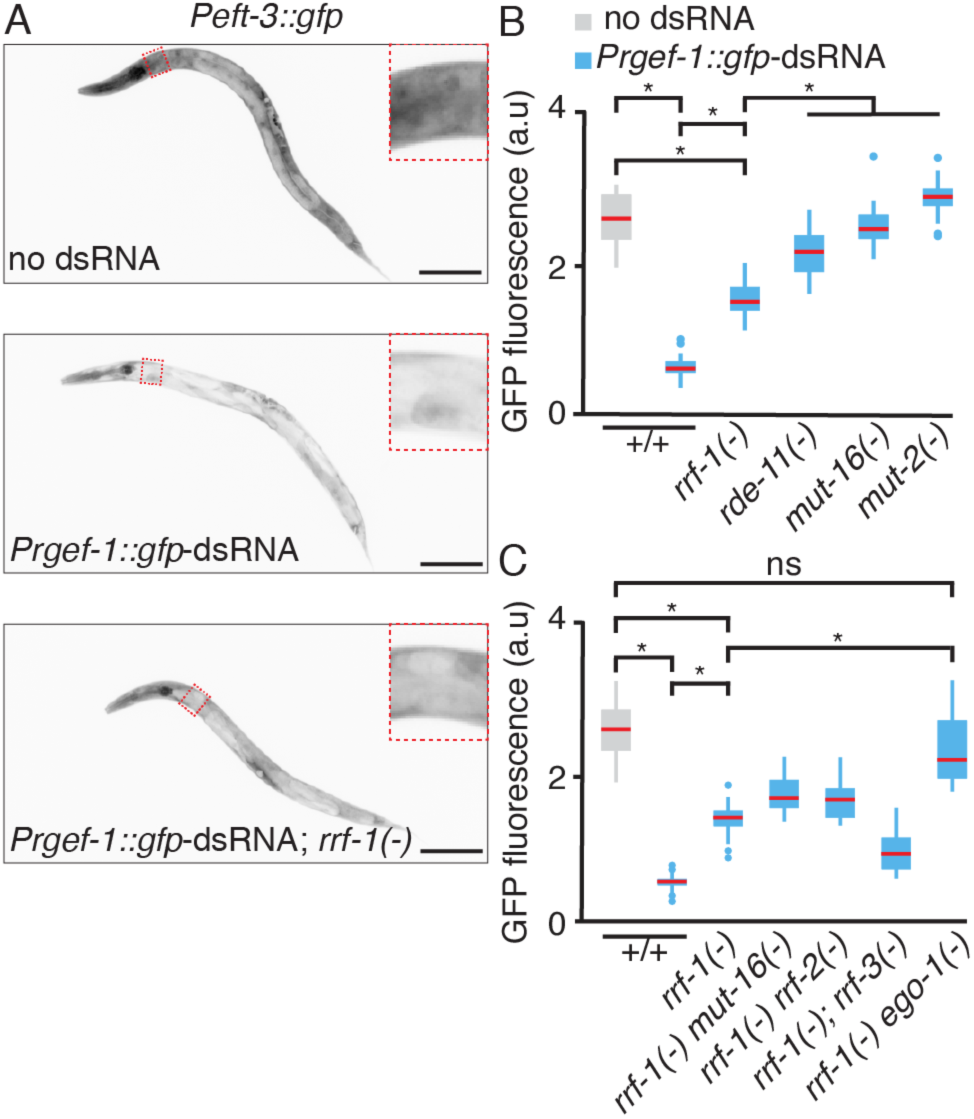
Silencing that can bypass a requirement for RRF-1 requires EGO-1 and MUT-16. (**A**) Silencing by neuronal dsRNA in the absence of RRF-1 is detectable for single-copy target sequences. Representative L4-staged animals that express GFP from a single-copy transgene in all tissues (*Peft-3::gfp*, *top*) and animals that in addition express *Prgef-1::gfp-*dsRNA in *rrf-1(+)* or *rrf-1(-)* backgrounds (*middle* or *bottom*, respectively) are shown. Insets are representative of the region quantified in multiple animals in B. Scale bar = 50 µm. Also see Supplementary Figure S3 for additional targets. (**B**) Silencing of *Peft-3::gfp* in the absence of *rrf-1* requires *rde-11*, *mut-16*, and *mut-2/rde-3.* GFP fluorescence was quantified (using arbitrary units (a.u.) in regions illustrated in (A) in control animals that do not express *Prgef-1::gfp-*dsRNA (grey) and in animals that express *Prgef-1::gfp-*dsRNA (blue) in wild-type (+/+), *rrf-1(-)*, *rde-11(-)*, *mut-16(-)* or *mut-2(-)* backgrounds. (**C**). The RdRP EGO-1 is required for silencing *Peft-3::gfp* in the absence of RRF-1, while the putative RdRP RRF-2 and the known RdRP RRF-3, do not compensate for the absence of RRF-1. As in (B), GFP fluorescence was quantified in control animals that do not express *Prgef-1::gfp-*dsRNA (grey) and in animals that express *Prgef-1::gfp-*dsRNA (blue) in wild-type (+/+), *rrf-1(-)*, *rrf-1(-) mut-16(-)*, *rrf-1(-) rrf-2(-)*, *rrf-1(-); rrf-3(-)*, *or rrf-1(-) ego-1(-)* backgrounds. Red bars indicate medians, asterisks indicate p-value < 0.05 (Student’s t-test) and n > 25 L4-staged animals except in *rrf-1(-) ego-1(-)* where n = 11. See Supplementary Figure S4 for details of *rrf-2*, *rrf-3* and *ego-1* alleles.

**Figure 3:**
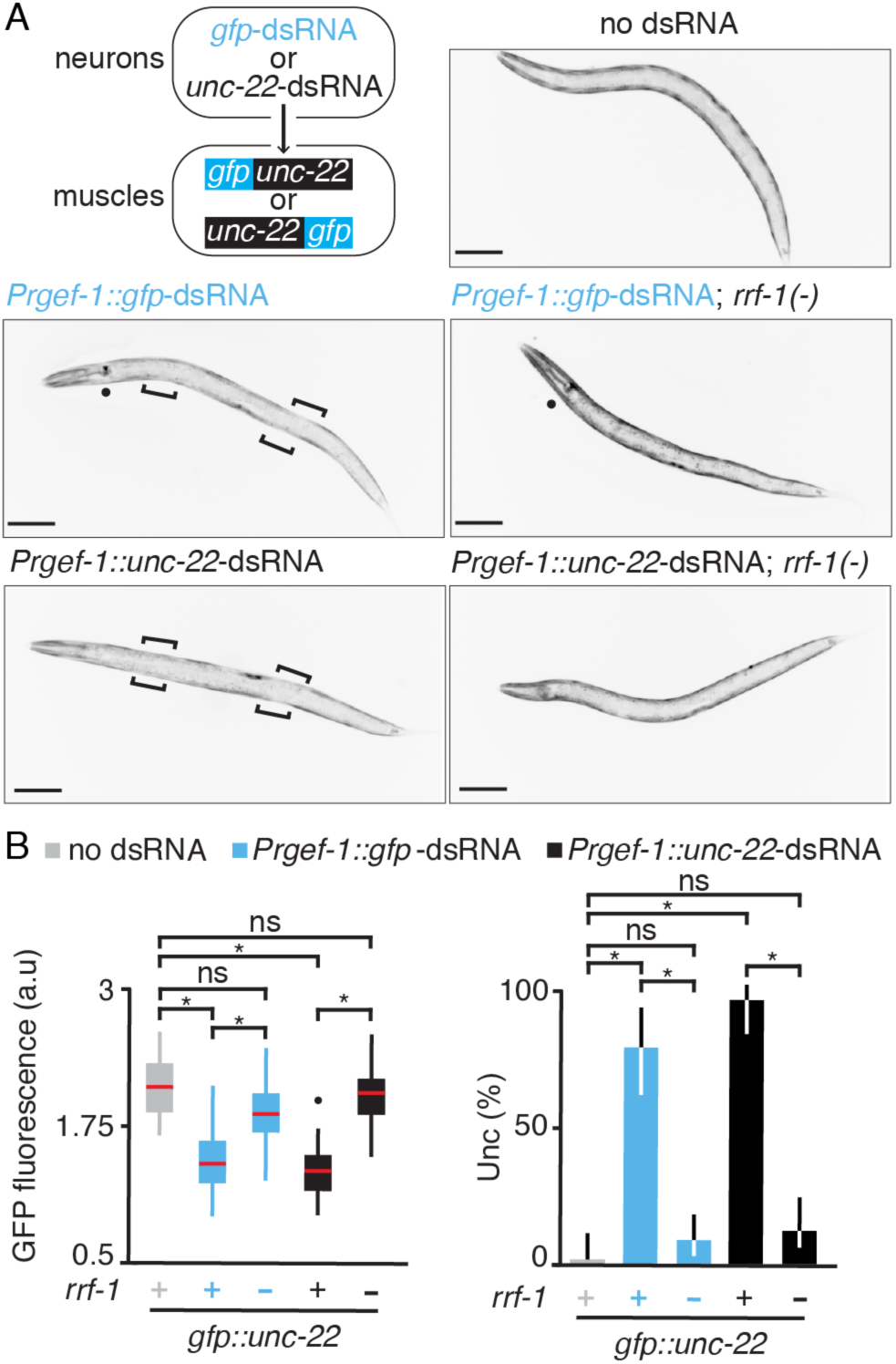
Changing the gene context of a target sequence can change the RRF-1 requirement for silencing that sequence. (**A**) *Top left*, Strategy for combining target sequences from experiments that showed different RRF-1 requirements to test silencing of a single chimeric target by neuronal dsRNA. The *gfp* sequence (blue) was inserted into the *unc-22* gene (black) at either the 5’ or 3’ ends to generate single chimeric target genes that can be silenced by either *gfp*-dsRNA or *unc-22*-dsRNA. See Supplementary Figure S4 for details of *gfp* insertions. *Top right*, Representative L4-staged animals that express GFP from *Punc-22::gfp::unc-22* and animals that in addition express *Prgef-1::gfp-*dsRNA (blue*, middle*) or *Prgef-1::unc-22*-dsRNA (black*, bottom*) in *rrf-1(+)* (*left*) or *rrf-1(-)* (*right*) backgrounds are shown. Fluorescence in the pharynx is observed in cases where *Prgef-1::gfp*-dsRNA is present (*middle*) because of expression from the co-injection marker *Pmyo-2::DsRed2* (circle) detected through the filters for GFP (see Materials and Methods). Animals with *Prgef-1::unc-22*-dsRNA have *Prgef-1::DsRed2* as a co-injection marker, which results in similarly detectable bleedthrough signal in the head region (*bottom*). Brackets indicate regions of silencing. Scale bar = 50 µm. (**B**) *Left,* GFP fluorescence from the chimeric gene (*Punc-22::gfp::unc-22*) was quantified (posterior to the pharynx) in control animals (*rrf-1(+)*) that do not express dsRNA (grey) and in animals that express either *Prgef-1::gfp*-dsRNA (blue) or *Prgef-1::unc-22*-dsRNA (black) in *rrf-1(+)* or *rrf-1(-)* backgrounds. Red bars, a.u., and n are as in 2B, asterisks indicate p-value <0.05 (Student’s t-test), and ns = not significant. *Right,* Percentage of animals that showed twitching (%Unc) expected upon silencing *Punc-22::gfp::unc-22* was scored for all strains shown in (A). Error bars indicate 95% confidence intervals, asterisks indicate p-value < 0.05 (Student’s t-test), ns = not significant, and n=50 L4-staged animals. Also see Supplementary Figure S6 for silencing of another chimeric target, *Punc-22::unc-22::gfp*.

**Figure 4:**
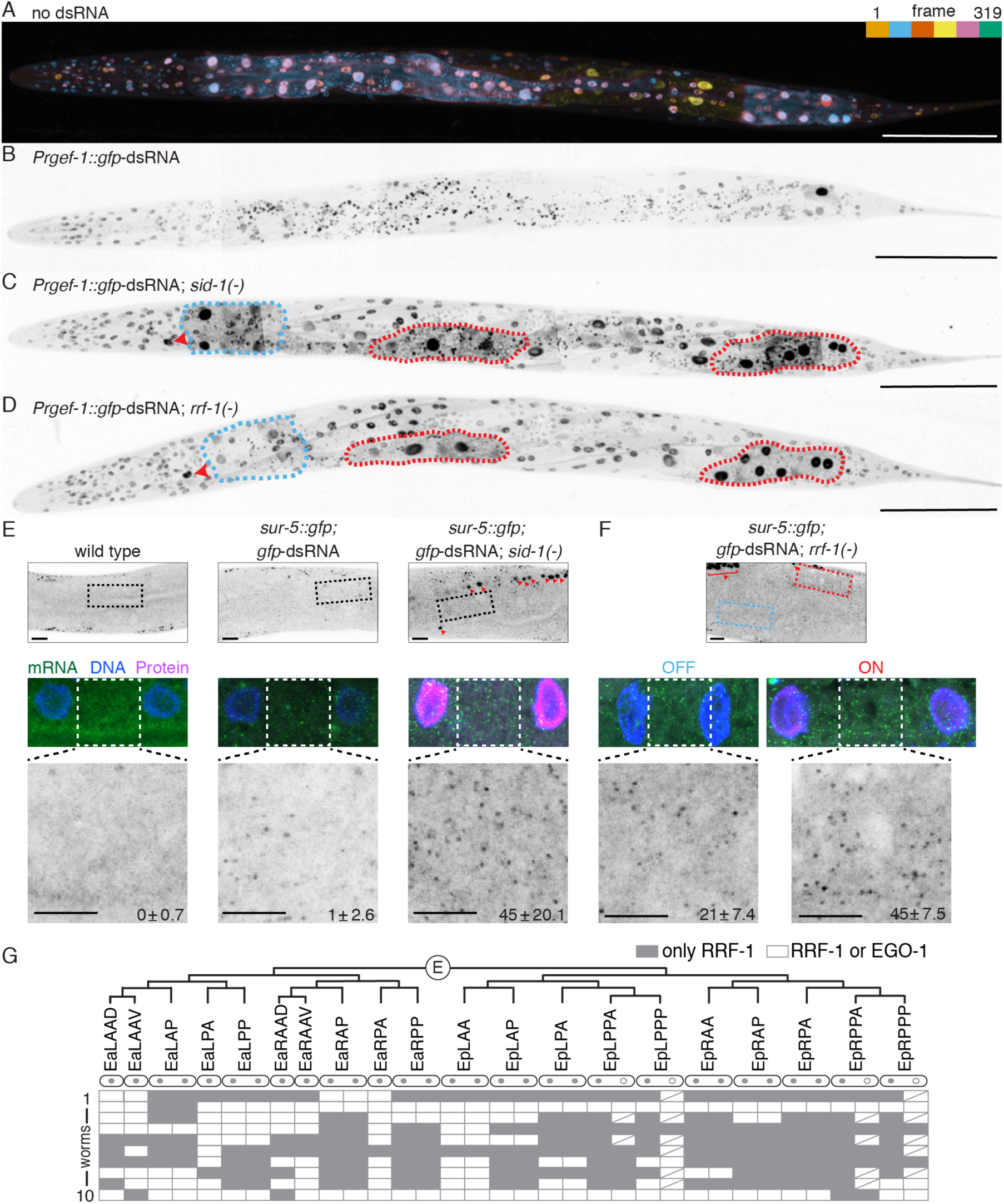
Identities of cells that require RRF-1 for silencing by neuronal dsRNA vary from animal to animal. (**A**) GFP expression from the *sur-5::gfp* chimeric gene enables simultaneous visualization of most somatic nuclei in *C. elegans*. A depth coded (one color for ∼53 frames) projection of 5 Z-stacks that were stitched together from a single L4-staged animal is shown (also see Materials and Methods). Scale bar = 100 µm. (**B-D**) Expression of *gfp*-dsRNA in neurons causes silencing throughout the animal that is entirely dependent on SID-1 and partially dependent on RRF-1. Representative images of L4-staged *sur-5::gfp* animals that express *Prgef-1::gfp*-dsRNA (B) and additionally lack *sid-1* (C) or *rrf-1* (D) are shown. Maximum intensity projections of sections were stitched together to generate whole-worm images. Presence of *gfp-*dsRNA causes worms to twist because of the *rol-6* co-injection marker. Cells that require RRF-1 for silencing (e.g. the excretory canal cell indicated by red arrows and some intestinal cells in red dashed lines) and cells that can silence in the absence of RRF-1 (e.g. some intestinal cells shown in blue dashed lines) are highlighted in *Prgef-1::gfp-*dsRNA*; rrf-1(-)* and in *Prgef-1::gfp*-dsRNA; *sid-1(-)* animals. Scale bar = 100µm. **(E-F)** Silencing in *rrf-1(-*) animals by neuronal dsRNA is associated with a decrease in *sur-5::gfp* mRNA levels. Single molecule FISH was used to detect *sur-5::gfp* mRNA in L4-staged wild-type animals (E, *left*) or in *sur-5::gfp* animals that express *Prgef-1::gfp-*dsRNA (E, *middle*) and that in addition lack *sid-1* (E, *right*) or *rrf-1* (F). RNA from *Prgef-1::gfp-*dsRNA is prominently detected by *gfp* probes in neuronal nuclei (red arrows and bracket). A representative pair of intestinal nuclei is shown for each animal as an overlay of DNA (DAPI in blue), mRNA (*gfp* in green) and protein (GFP in magenta). Cytoplasmic mRNA foci were counted (see Materials and Methods) between two nuclei in wild-type or in *sid-1(-)* backgrounds (E), and between two nuclei where GFP is silenced (off, blue) and where GFP is expressed (on, red) in *rrf-1(-)* animals. Errors indicate 95% confidence intervals, n=3 in E and n=4 in F. Top scale bar = 10 µm and bottom scale bar = 5 µm. (**G**) Identities of intestinal cells that require RRF-1 for silencing vary from animal to animal. The E blastomere divides to generate 20 intestinal cells (EaLAAD to EpRPPP). Of the 20 cells, 10 undergo nuclear division without cell division (two grey circles per cell), 4 sometimes undergo similar nuclear division (one grey circle and one open circle per cell), and 6 do not undergo any division (one grey circle per cell). In each of 10 *sur-5::gfp; rrf-1(-); Prgef-1::gfp*-dsRNA L4-staged animals, GFP-positive nuclei (use only RRF-1, grey) and GFP-negative nuclei (use EGO-1 or RRF-1, white) were scored. Every binucleate cell had both nuclei of the same class. White boxes with a slash indicate absence of second nucleus because of lack of nuclear division (47). See Supplementary Figure S7 for images of additional animals.

### Image processing

All images being compared in a figure were adjusted identically using Adobe Photoshop and/or FIJI (NIH). Images taken on Nikon AZ100 were inverted (GFP = black), look-up tables were changed using Photoshop (190 = white and 255 = black for *gtbp-1::gfp*, *eft-3::gfp*, *gfp::unc-22* and *unc-22::gfp*; 212 = white and 255 = black for *sur-5::gfp*), and cropped for display. When imaging using the SP5X confocal (Figure 4A-D and Supplementary Figure S8), our immobilization conditions resulted in the worm lying on the coverslip such that the middle of the worm (vulva region) was tightly sandwiched between the coverslip and the agarose pad but the rest of the worm (head and tail in particular) was free to assume different positions. To partially account for this variability and the observed loss in sensitivity with depth of imaging, stacks close to the coverslip that lacked any signal were removed (0-30 stacks, median 7 stacks) and an equivalent number of empty stacks were added beyond the worm for a consistent total of 319 stacks in all cases. For Figure 4A-4D, Z-projections of the 5 stacks for each worm were stitched together using a combination of a pairwise stitching plugin (45) and manual alignment (Adobe Illustrator). For Figure 4A, each Z-stack was depth-coded using the ‘temporal-color code’ function in FIJI (6 colors with 53 stacks/color). For Supplementary Figure S8A, Z projections of maximum intensity were created using all 319 stacks (head and tail) or a subset of stacks (seam, uterus and vulva). For Figure 4E and 4F, Z projections of maximum intensity were created using 5 slices, inverted (GFP = black), cropped for display (full anterior region or zoomed-in region between two nuclei) and look-up tables were changed using Photoshop (160 = white and 255 = black). Composites of GFP, DAPI and Quasar 670 were created on FIJI (NIH) and look-up tables were changed to magenta, blue, or green.

### Quantification of silencing

Silencing in response to *unc-22-*dsRNA was scored by calculating the percentage of L4-staged animals that twitched within 3 min in 3mM levamisole. The silencing of GFP expressed from *nrIs20* (*sur-5::gfp)* was determined by counting the number of intestinal nuclei that showed bright GFP fluorescence in L4-staged animals at a fixed magnification and zoom using a MVX10 stereomicroscope (Olympus). Average number of intestinal nuclei were determined by counting HC195 and was relatively constant in most genetic backgrounds with the exception of strains that lacked *rrf-1* (e.g. 32.8 ± 0.6 nuclei in *rrf-1(-); nrIs20* animals and 32.3 ± 0.8 nuclei in *rrf-1(-); eri-1(-) nrIs20* animals, compared to 29.9 ± 1.2 nuclei in *nrIs20* animals, errors indicate 95% CI). For images acquired using Nikon AZ100, silencing was quantified using FIJI (NIH) by measuring the fluorescence posterior to the pharynx in a region of interest (ROI) that included either a fixed area anterior to the germline (Figure 2 and Supplementary Figure S3) or body-wall muscles all along the worm (Figure 3 and Supplementary Figure S6), using the formula ROI fluorescence (arbitrary units) = intensity of ROI – (area of ROI x mean intensity of background). For images acquired using the SP5X confocal microscope, a combination of thresholding using the 3D object counter plugin (46) on FIJI (NIH) and manual verification was used to count various nuclei. To score nuclei as ‘on’ or ‘off’, different thresholds were used for intestinal nuclei located at different depths (70 for stacks 1-160; 20 for stacks 161-319) and a constant threshold was used for all other nuclei (20 for all stacks). For Figure 4E and F, the number of mRNA foci was counted using the 3D object counter on FIJI (NIH). A threshold of 50 was selected, objects <0.015 µm^3^ were eliminated as background, and objects >0.2 µm^3^ were eliminated as miscounts due to merging of multiple objects. For Figure 4G, the identity of each intestinal nucleus was inferred using its expected location and using the position of the vulva, anus, and the twisting rows of hypodermal cells (twist induced by the *rol-6* co-injection marker for *gfp-*dsRNA [qtIs49]) as guideposts (47–52).

### Statistics

Significance of differences in silencing (p-value < 0.05, unless otherwise stated) were calculated using Student’s t-test (two tailed) or a two-way Analysis of Variation (ANOVA) with replication. Error bars in Figure 3B, *Right* and Supplementary Figure S6A and S6C, *Right* indicate 95% confidence intervals for single proportions calculated using Wilson’s estimates with a continuity correction (Method 4 in (53)) and significance of differences between strains was determined using pooled Wilson’s estimates.

## RESULTS

### Silencing by neuronal dsRNA can be distinct from silencing by ingested or target-derived dsRNA

Double-stranded RNA can be introduced into *C. elegans* cells through the transcription of complementary sequences within the target cell, in a distant cell, or in ingested bacteria. While all these sources of dsRNA trigger RDE-1-dependent gene silencing (54), each source could produce different pools of dsRNA and/or dsRNA-derivatives that are trafficked differently to the cytosol of the target cell where silencing occurs. Here we present evidence that different sources of dsRNA can differ in their requirement for RRF-1 to silence the same target gene.

To examine silencing of a single target by different sources of dsRNA, we used a nuclear-localized GFP that is expressed in all somatic cells (*sur-5::gfp*) and is particularly prominent in the large intestinal nuclei (Figure 1A, *Top left*, ∼30 GFP+ nuclei). This target is a multicopy transgene that generates trace amounts of dsRNA that can cause self-silencing in enhanced RNAi backgrounds (e.g. *adr-1(-); adr-2(-)* in (55) and *eri-1(-)* or *rrf-3(-)* in (30)). Silencing by this target-derived dsRNA was modest (Figure 1A, ∼24 GFP+ nuclei in *eri-1(-)*, p-value *<* 10^-3^ when compared to ∼30 GFP+ nuclei in *eri-1(+)*), consistent with earlier reports (11, 30). Similarly, silencing by *gfp-*dsRNA expressed in neurons (*Prgef-1::gfp*-dsRNA) was also modest (Figure 1A, ∼24 GFP+ nuclei, p-value *<* 10^-4^ when compared to *eri-1(+)*), consistent with an earlier report (12). However, when both target-derived and neuronal dsRNA were present together (i.e. in *eri-1(-); Prgef-1::gfp*-dsRNA animals), we observed a synergistic effect resulting in greatly enhanced silencing (Figure 1A, ∼3 GFP+ nuclei, two-way ANOVA p-value < 10^-20^ for interaction). This enhancement, taken together with the previous observation that ERI-1 inhibits silencing by neuronal *unc-22*-dsRNA (Supplementary Figure 3 in ref. (12)), suggests that ERI-1 inhibits silencing by *gfp*-dsRNA generated from the target and *gfp*-dsRNA imported from neurons (Figure 1B). Upon performing a genetic screen using these robustly silenced animals, we isolated alleles of four genes with known roles in RNAi - *rde-1*, *rde-11*, *sid-1*, and *rrf-1* (Figure 1C, Supplementary Figure S1). Surprisingly, unlike in null mutants of *rde-1, rde-11*, or *sid-1*, significant silencing (p-value < 10^-7^) was detectable in null mutants of *rrf-1* (Figure 1C) – a property shared by all three alleles of *rrf-1* isolated in the screen (Figure 1C). Tissue-specific rescue experiments suggest that both *rde-1* and *rrf-1* function in the intestine (target cells) and not in neurons (source cells) to enable the observed silencing of intestinal cells (Supplementary Figure S2). Thus, when both target-derived dsRNA and neuronal dsRNA were used together to silence the same gene, RDE-1-dependent but RRF-1-independent silencing was detectable in some intestinal cells.

This bypass of RRF-1 could be a feature of silencing by target-derived dsRNA, neuronal dsRNA, or a general feature of silencing by all sources of dsRNA. To determine RRF-1 requirements for silencing by different sources of dsRNA, we examined silencing by target-derived dsRNA using an *eri-1(-)* background, silencing by neuronal dsRNA in an *eri-1(+)* background, and silencing by ingested dsRNA. All three sources of dsRNA strictly required RDE-11, a dosage-sensitive RNAi factor (56, 57). In contrast, the requirement for RRF-1 varied depending on the source of dsRNA. The weak silencing by target-derived dsRNA was completely abolished in *rrf-1* null mutants (Figure 1D orange). Equally weak silencing by neuronal dsRNA was not significantly altered in *rrf-1* null mutants (Figure 1D blue). Yet, robust silencing by ingested dsRNA was strongly dependent on RRF-1 (Figure 1D black). These source-dependent differences in extents of silencing could be caused by differences in the routes taken by dsRNA to reach the silencing machinery, the forms of dsRNA and/or the dosages of dsRNA. However, because weak silencing by neuronal dsRNA was partially independent of RRF-1, while strong silencing by ingested dsRNA was primarily dependent on RRF-1, a high dose of dsRNA from neurons cannot be the sole explanation for the observed RRF-1 independence. Therefore, these observations suggest that mechanisms engaged by ingested or target-derived dsRNA can differ from those engaged by neuronal dsRNA.

### EGO-1 can compensate for lack of RRF-1

To determine if other loci could show silencing by neuronal dsRNA in the absence of RRF-1, we used the same source of neuronal dsRNA and examined silencing of GFP expression under the control of a different promoter introduced into different genomic loci. Silencing of *gfp* expressed under the control of the *eft-3* promoter (*Peft-3::gfp*) from a single-copy transgene was partially independent of RRF-1 (Figure 2A). In the absence of RRF-1, a significant reduction in GFP fluorescence was detectable (Figure 2B). A similar extent of silencing in *rrf-1(-)* animals was observed using *Peft-3::gfp* transgenes located on three different chromosomes (Supplementary Figure S3A) and for a C-terminal *gfp* fusion of a ubiquitously expressed gene (Supplementary Figure S3B and Supplementary Figure S3C) generated using Cas9-based genome editing (Supplementary Figure S4). Thus, a measurable amount of silencing by neuronal dsRNA can occur in the absence of RRF-1 when *gfp* is expressed under the control of different promoters and from different chromosomes.

Although it is formally possible that neuronal dsRNA engages novel processing pathways that are not used by other sources of dsRNA, we found that additional components of canonical RNAi were required for silencing (Figure 2B and Supplementary Figure S5). RDE-11, thought to facilitate the production of secondary siRNA (56, 57), was required for most silencing (Figure 2B). MUT-16, a poly-Q/N protein (58) and MUT-2/RDE-3, a putative nucleotidyltransferase (59), that together localize to perinuclear foci thought to be sites of secondary siRNA production (60, 61), were both required for all observed silencing (Figure 2B). Consistently, GFP protein levels in *mut-16(-)* animals were greater than that in *rrf-1(-)* animals (Supplementary Figure S3D). Removal of MUT-16 in the *rrf-1(-)* background (Supplementary Figure S4) resulted in weaker silencing of this target (see persistent nuclear fluorescence in Supplementary Figure S5) and complete loss of silencing for another target (see below). These results suggest that silencing by neuronal dsRNA in the absence of RRF-1 either occurs through the action of primary siRNAs along with canonical factors such as RDE-11, MUT-16, and MUT-2/RDE-3, or through the production of secondary siRNAs using an alternative RdRP.

The *C. elegans* genome has four genes that encode proteins with RdRP domains, three of which have demonstrated roles in the production of RNA using RNA templates. RRF-3 is thought to act as a processive RdRP in an endogenous pathway (62, Supplementary Figure 9 in (63)) that competes with experimental RNAi for shared components (64) and therefore loss of *rrf-3* enhances RNAi (65). RRF-1 and EGO-1 are thought to act as non-processive RdRPs that make siRNAs in the soma (20, 21, 64) and the germline (66), respectively. Preventing germline proliferation in *rrf-1(-)* animals was found to greatly reduce the levels of secondary siRNAs but not eliminate them (67), leaving open the possibility that the residual secondary siRNAs may be generated by an alternative RdRP. The fourth putative RdRP, RRF-2, was found to be not required for silencing by ingested dsRNA (22). To test if the silencing observed in the absence of RRF-1 depends on any of these other RdRPs, we generated mutants lacking RRF-2, RRF-3, or EGO-1 using Cas9-based genome editing (Supplementary Figure S4). In an *rrf-1(-)* background, loss of *rrf-2* did not eliminate silencing and loss of *rrf-3* resulted in enhancement of silencing (Figure 2C and Supplementary Figure S5). Evaluation of the loss of *ego-1* is complicated by the sterility of *ego-1(-)* animals, reflecting the role of EGO-1 in germline development (23, 24). However, *ego-1(-)* progeny of heterozygous animals lacked all silencing in the absence of *rrf-1* despite the potential for parental rescue of *ego-1* (Figure 2C and Supplementary Figure S5), suggesting that EGO-1 made in progeny compensates for the absence of RRF-1. Hereafter, we shall refer to silencing in the absence of RRF-1 as silencing using EGO-1.

Taken together, these results reveal instances of silencing in somatic cells by a source of neuronal dsRNA through the use of two different RdRPs.

### Context of target sequence can dictate RdRP usage

Expression of dsRNA in neurons does not always cause detectable silencing in the absence of RRF-1, suggesting that EGO-1 is not used in all contexts. For example, neuronal dsRNA targeting *unc-22* ((12), Supplementary Figure S6A) or *bli-1* (Supplementary Figure S6A) required RRF-1 for all silencing. Nevertheless, targeting *gfp* sequences using neuronal dsRNA resulted in silencing using EGO-1 in animals that lack *rrf-1* (in Figure 1, Figure 2, and Supplementary Figure S3 using an integrated *gfp*-dsRNA source, and in 6/6 *rrf-1(-); gtbp-1::gfp* animals using an extrachromosomal *gfp*-dsRNA source). These results suggest that silencing in somatic cells using EGO-1 is not a generic property of all neuronal dsRNA and raise two possibilities: (1) sources that do not strictly require RRF-1 (e.g. neuronal *gfp*-dsRNA) differ from sources that require RRF-1 (e.g. neuronal *unc-22*-dsRNA); or (2) target sequences that do not strictly require RRF-1 (e.g. *gfp*) differ from target sequences that require RRF-1 (e.g. *unc-22*).

To examine silencing of a single target sequence by either source of dsRNA, we generated two chimeric genes (*gfp::unc-22* or *unc-22::gfp*) that could both be silenced by either *gfp*-dsRNA or *unc-22*-dsRNA expressed in neurons (Figure 3A, *top left*). Both chimeric genes express *unc-22* and *gfp* sequences as a single transcript under the control of endogenous *unc-22* regulatory sequences (Supplementary Figure S4) and were functional as evidenced by lack of twitching (Figure 3B, *right*), which is a sensitive readout of reduction in *unc-22* function (2). With either source of dsRNA, all measurable silencing required RRF-1 (Figure 3, Supplementary Figures 6B and 6C). This complete dependence on RRF-1 was more evident when twitching was measured in response to the expression of either *gfp-*dsRNA or *unc-22-*dsRNA in neurons (Figure 3B, *right*).

These results suggest that changing the context of a target sequence can change its need for RRF-1 versus the alternative use of EGO-1 for silencing by neuronal dsRNA. Specifically, silencing of the single-copy *gfp* target by neuronal *gfp*-dsRNA could use EGO-1 when *gfp* is present as part of *eft-3::gfp* or *gtbp-1::gfp* but not as part of *unc-22::gfp*. These differences in genomic location, associated regulatory elements, or site of expression could be responsible for the observed differential use of EGO-1.

### Somatic cells that can use EGO-1 for silencing vary from animal to animal

To examine the use of EGO-1 for silencing in all somatic cells while keeping the genomic location and associated regulatory elements of the target gene constant, we generated a chimeric gene with *gfp* sequence fused to the endogenous *sur-5* gene (*sur-5::gfp,* Supplementary Figure S4). This strain resulted in the expression of a nuclear-localized SUR-5::GFP fusion protein, enabling simultaneous visualization of every somatic nucleus using confocal microscopy (Figure 4A). Expression of *gfp*-dsRNA in neurons resulted in silencing throughout the length of the animal that was entirely dependent on SID-1, consistent with silencing by neuronal dsRNA (Figure 4B and Figure 4C) and was not subject to silencing upon *eri-1* loss by target-derived dsRNA (Supplementary Figure S7A-C) as is expected for a single-copy target (30). Silencing was easily detected in intestinal cells, hypodermal cells, body-wall muscle cells, and the excretory canal cell (Figure 4B and Supplementary Figure S8A). Silencing was not detectable in some cells in the head, the vulval and uterine regions, and occasionally in the tail region (Supplementary Figure S8A). Interestingly, even neighboring, lineal sister cells sometimes showed very different extents of silencing (e.g., intestinal cells near the tail in Supplementary Figure S8A, *top row*). Nevertheless, the overall silencing observed was much more than that observed when the same source of dsRNA was used to silence a multi-copy *sur-5::gfp* transgene (1.0±0.4 visible intestinal nuclei for single-copy *sur-5::gfp* (Supplementary Figure S7C) versus 24.0±1.9 visible intestinal nuclei for multi-copy *sur-5::gfp* (Figure 1A), p-value < 10^-21^ and errors indicate 95% CI). A simple explanation for this difference could be that silencing higher numbers of target transcripts requires higher amounts of dsRNA (see Discussion for additional possibilities). Thus, the single-copy *sur-5::gfp* gene is a sensitive target for evaluating the use of EGO-1 for silencing by neuronal dsRNA in somatic cells throughout the animal.

Silencing of single-copy *sur-5::gfp* by neuronal dsRNA was detectable in *rrf-1(-)* animals (Figure 4D), but the extent of silencing and the locations of cells that showed silencing varied dramatically from animal to animal (Supplementary Figure S7D). To obtain a high-resolution view of silencing, we quantified silencing in multiple tissues by counting the number of nuclei that show fluorescence (Supplementary Movie S1). For quantifying silencing in hypodermal and body-wall muscle cells, we divided the body into three regions (Supplementary Figure 8B, *Left*): head (anterior to the posterior bulb of the pharynx), anterior body (anterior to the vulva), and posterior body (posterior to the vulva). In the head and anterior body, the average numbers of detectable nuclei in *rrf-1(-)* animals were not very different from the average numbers detectable in *sid-1(-)* animals (Supplementary Figure S8B, *Right*). The posterior body, however, showed marginal silencing of hypodermal and/or body-wall muscle cells in *rrf-1(-)* animals (50.0±7.6 nuclei versus 58.7±4.6 nuclei in *sid-1(-)*, p-value = 0.08 and errors indicate 95%CI), suggestive of some use of EGO-1 for silencing. The intestine, however, showed obvious silencing in the absence of RRF-1. This silencing was associated with a reduction in mRNA levels (Figure 4E and 4F) and required MUT-16 and EGO-1 (Supplementary Figure 7E). Notably, loss of EGO-1 alone does not result in a detectable defect in silencing by neuronal dsRNA (Supplementary Figure 7F), suggesting that EGO-1 is not required for silencing in any intestinal cell but rather can compensate for loss of RRF-1.

Because each of the 20 intestinal cells has an invariable lineal origin and position after morphogenesis (Figure 4G, (47–52)), we were able to examine whether silencing occurs in any discernable patterns correlated with lineage or position. Each tested worm had a different complement of cells with respect to RdRP use for silencing (Figure 4G and Supplementary Figure S7D) such that no cell relied on only RRF-1 in every animal and no cell could use EGO-1 in every animal (Figure 4G).

Together, these results show that neuronal dsRNA can cause robust silencing, but the particular cells that require RRF-1 for such silencing vary from animal to animal.

## DISCUSSION

We examined RNA interference in the somatic cells of *C. elegans* and found that the source of extracellular dsRNA, the context of target sequences, and the identity of the tested cell can all dictate whether the RNA-dependent RNA polymerase RRF-1 is required for silencing. We discovered that silencing by neuronal dsRNA can be extensive and, when examined at single-cell resolution, different sets of cells rely on only RRF-1 or could also use EGO-1 in the absence of RRF-1 for silencing in each animal.

### Silencing by neuronal dsRNA

Expression of dsRNA in all neuronal cells resulted in SID-1-dependent silencing in a variety of cell types throughout the animal (hypodermal cells, body-wall muscle cells, seam cells, intestinal cells, and excretory canal cell; Figure 4 and Supplementary Figure S8), suggesting that dsRNA molecules exported from neurons are widely available. Subsequent import depends on the levels of SID-1 in importing cells because cells that overexpress SID-1 can act as sinks for dsRNA and presumably reduce entry of dsRNA into other cells (Supplementary Figure S2 in (11), (68)). The observed widespread silencing (Figure 4) therefore suggests that no single tissue acts as a sink and that sufficient dsRNA is exported from neurons to reach cells throughout the animal.

Yet, silencing by neuronal dsRNA is not always detectable in all cells, which could reflect either inefficient import of dsRNA or inefficient silencing. For example, most intestinal cells were not silenced when neuronal dsRNA was used to silence a multi-copy *sur-5::gfp* transgene (Figure 1A). However, silencing of this multi-copy target was greatly enhanced upon loss of *eri-1* (Figure 1A), which releases shared factors used for endogenous RNAi-related processes (69). Therefore, this case of limited silencing by neuronal dsRNA likely reflects limited availability of such RNAi factors (e.g. RDE-4, DCR-1, etc.) and not poor access to dsRNA or poor import of dsRNA. Similarly, the lack of silencing of single-copy *sur-5::gfp* in the cells of some tissues (pharynx, vulva, and uterus, Supplementary Figure S8A) could reflect inefficient silencing that could potentially be enhanced by providing limiting factors.

### Cellular origins of small RNAs

A wide range of endogenous small RNAs (miRNAs, siRNAs, piRNAs etc.) are being analyzed by sequencing RNA from whole worms. Where any particular small RNA is made and where it acts are both obscured when worms are homogenized for extracting RNA. Base-paired RNAs such as long dsRNA (70), precursors of miRNAs (17, 18) or precursors of 26G RNAs (63, 64) could be transported through SID-1 such that they are made in one cell and cause effects in other cells. However, tests for such non-autonomous effects of the *lin-4* miRNA suggest cell-autonomous action of this miRNA (71). Examination of some of the numerous anti-sense RNAs called 22G RNAs suggested that they are made by RRF-1 in somatic cells and both RRF-1 and EGO-1 in the germline (72). Our results suggest the possibility that some 22G RNAs could be made in the intestine in the absence of RRF-1 potentially using EGO-1 in the intestine or through indirect effects of EGO-1 function in the germline. Resolving the origin and the site of action of such endogenous small RNAs requires controlled experiments that consider both non-cell autonomy of the RNA and functional mosaicism of its biogenesis.

### Functional mosaicism of RNAi in an animal

The identities of the intestinal cells that strictly require RRF-1 for silencing by neuronal dsRNA varied from animal to animal (Figure 4G and Supplementary Figure S7). This variation observed in *rrf-1(-)* animals could be because of unequal and random availability of compensatory EGO-1 despite equal availability of neuronal dsRNA or because of unequal and random availability of neuronal dsRNA despite equal availability of compensatory EGO-1 (Supplementary Figure S9). Such functional mosaicism is masked in wild-type animals, where the amplification of silencing signals by RRF-1 results in uniform silencing. Thus, RRF-1 promotes silencing by extracellular dsRNA to ensure uniform silencing - a role that is reminiscent of the role for ERI-1 in opposing silencing by transgene-derived dsRNA to ensure uniform expression (30).

RNAi is an antiviral mechanism in many organisms (see (73) for a recent evolutionary perspective) and wild strains of *C. elegans* that are defective in RNAi can harbor viruses (74). Viral infection of *C. elegans* in the lab results in proliferation of the virus in some but not all intestinal cells (75). It would be interesting to determine whether mosaicism of specific components of the RNAi machinery underlies patterns of viral infection observed in the intestine of *Caenorhabditis* nematodes (74, 75). Analyses of variation in intact animals where organismal regulatory mechanisms are preserved, as described here using *C. elegans,* are an effective complement to analyses in single cells, which have begun to reveal heterogeneity in many processes (e.g. in gene expression (76), in membrane trafficking (77), and in subcellular organization (78)). We speculate that functional mosaicism and its control could be common in multicellular organisms because of the need to balance diversification of cell types with preservation of fundamental functions in all cells.

## DATA AVAILABILITY

All strains and original confocal images are available upon request. Whole genome sequencing data for *rrf-1(jam2, jam3* and *jam4), rde-1(jam1), rde-11(jam50* and *jam51)* and *sid-1(jam52)* are available on Sequence Read Archive (PRJNA486008).

## ACKNOWLEDGEMENTS

We thank Leslie Pick and members of the Jose lab for critical comments on the manuscript; Amy Beaven at the University of Maryland’s imaging core for microscopy advice, Suwei Zhao at the IBBR sequencing facility for whole genome sequencing of mutants, Molly Lutrey for complementation analysis of *rde-11* mutants, Pravrutha Raman for preparation of genomic DNA from two mutants, the *Caenorhabditis elegans* Genetic stock Center, the Hunter lab (Harvard University), the Seydoux lab (Johns Hopkins University) and the Jorgensen lab (University of Utah) for some worm strains, and the Hamza lab (University of Maryland) for bacteria that express *gfp*-dsRNA.

## SUPPLEMENTARY DATA

### SUPPLEMENTARY FIGURES AND FIGURE LEGENDS

**Supplementary Figure S1:**
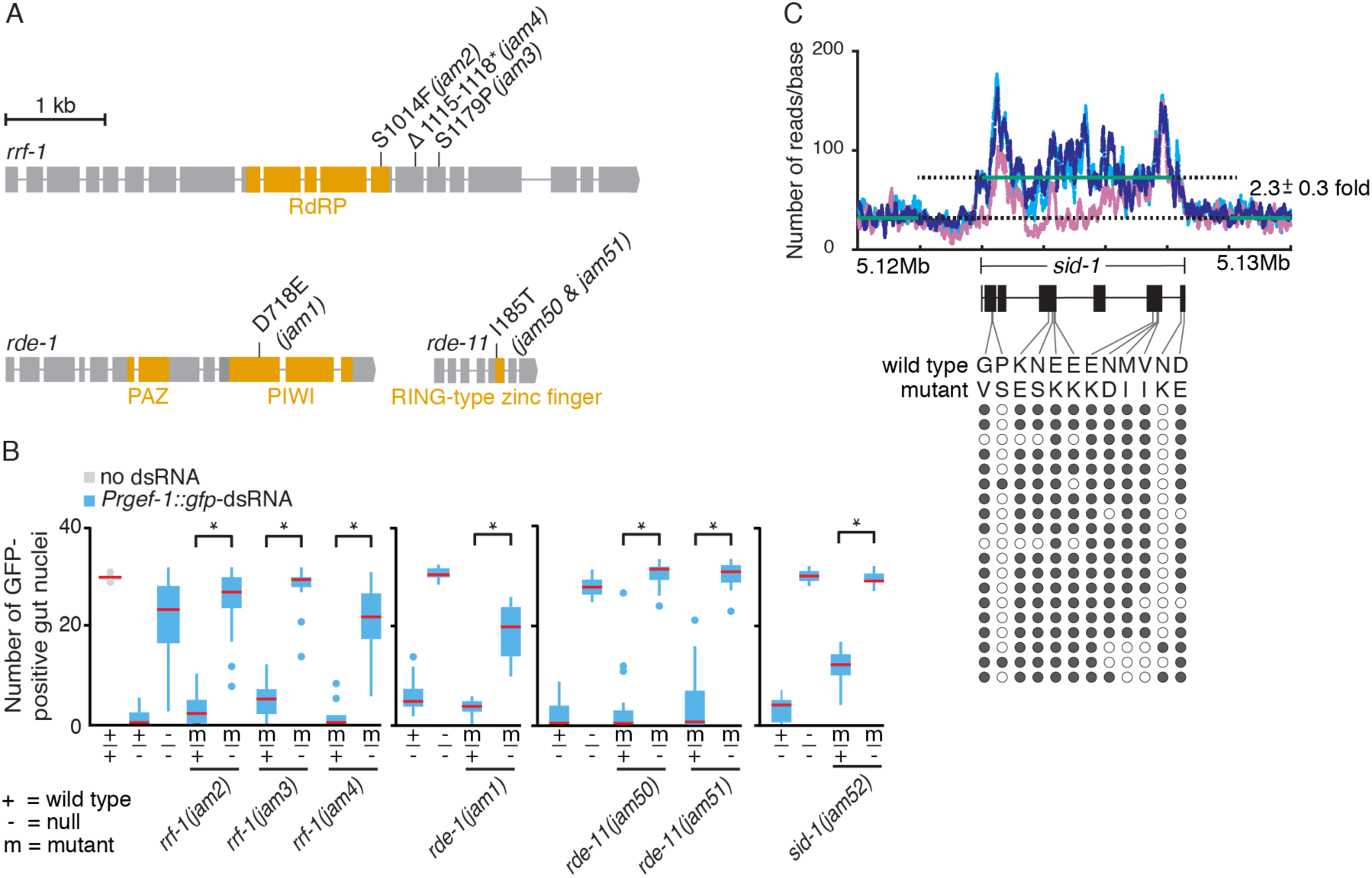
Four genes with known roles in RNAi were identified among mutants defective in silencing upon *eri-1* loss and/or by neuronal dsRNA. (A) Schematic of *rrf-1*, *rde-1*, and *rde-11* genes with identified molecular lesions. The gene structures (exons = boxes, introns = lines), identified alleles along with deduced changes in the corresponding proteins (*jam2*(S1014F), *jam3*(S1179P), and *jam4(Δ*1115-1118*, short insertion/deletion resulting in downstream stop codon) in *rrf-1*, *jam1*(D718E) in *rde-1*, and *jam50*(I185T) and *jam51*(I185T) in *rde-11*), and regions encoding known protein domains (in orange, RdRP in *rrf-1*, PAZ and PIWI in *rde-1*, and RING-type zinc finger in *rde-11*) are indicated. Scale bar = 1 kb. (B) Each isolated allele fails to complement its corresponding null allele. Silencing in test crosses was scored by counting the number of GFP-positive intestinal nuclei in male cross progeny (m/-) when mutant animals (m) were each crossed to animals with a null allele *(-).* Male progeny of mutants crossed with N2 (m/+) and of the null mutant crossed to the strain used for mutagenesis (AMJ1, see strain list for complete genotype) (+/-) were scored as controls. Red bar, n and asterisks are as in Figure 1C. Unlike in *rde-1* null mutants, silencing is detectable in *rde-1(jam1)* animals, indicating that the *jam1* mutation results in a partial loss of function. Both *jam50* and *jam51* mapped to chromosome IV and an examination of all mutations identified by whole genome sequencing revealed identical mutations in *rde-11* and in other genes, suggesting that these mutants are siblings. Sequencing of the exons of *sid-1* in animals with *jam52* did not reveal any mutations although *jam52* maps to chromosome V and complements *rde-1* but not *sid-1*. (C) One of the transgenes present in AMJ1 includes a mutated copy of *sid-1* gene sequence incorporated from a PCR fragment. Illumina sequencing reads covering each base for three different mutants (cyan, navy, magenta) indicate that the average coverage (green lines) within the *sid-1* gene is ∼2 fold higher than that in surrounding regions, consistent with the presence of one additional copy in the background. Twelve changes in the *sid-1* gene (in >15% of reads covering a base) that altered the encoded amino acid (wild-type and mutant residues are indicated) were detected in at least 2 of 19 sequenced mutants (filled circles). Consistently, this mutated copy of *sid-1*, which is a part of the *qtIs50* transgene, is non-functional (95.3% silencing of *dpy-7* in *qtIs50* animals versus 1.7% silencing in *sid-1(-); qtIs50* animals by feeding RNAi, n>40 L4-staged animals).

**Supplementary Figure S2:**
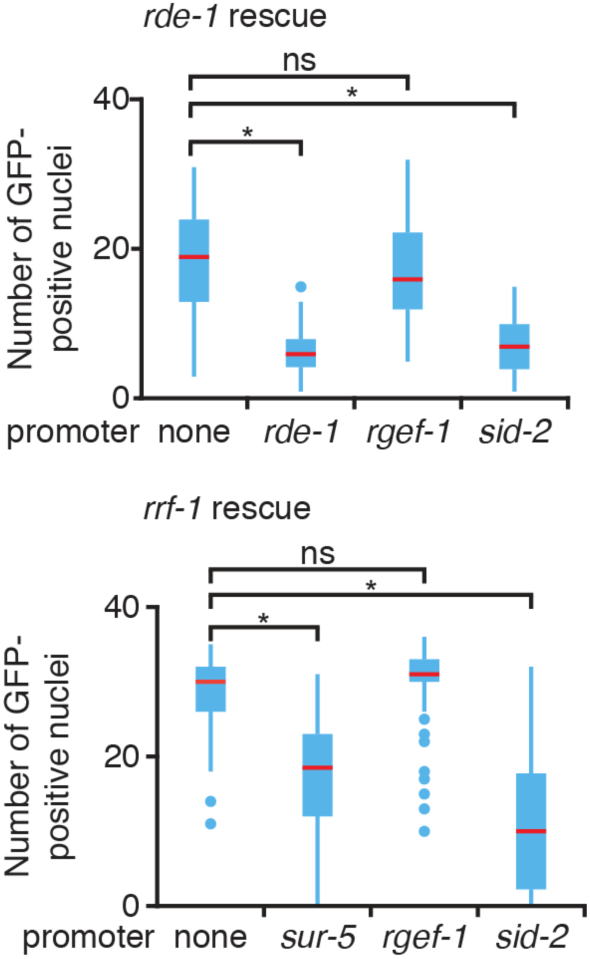
*rde-1* and *rrf-1* are required in the intestine for silencing by neuronal dsRNA. *Top*, Rescue of *rde-1(jam1)* using tissue-specific promoters indicates that RDE-1 functions in intestinal cells but not in neurons to enable silencing by neuronal dsRNA. The number of GFP-positive intestinal nuclei in *rde-1(jam1)* animals that were transformed with either a co-injection marker alone (none) or a co-injection marker along with *rde-1(+)* expressed under its own (*rde-1*), intestine-specific (*sid-2*), or neuron-specific (*rgef-1*) promoter were counted. *Bottom*, Rescue of *rrf-1(jam3)* using tissue-specific promoters indicates that RRF-1 functions in intestinal cells but not in neurons to enable silencing by neuronal dsRNA. The number of GFP-positive intestinal nuclei in *rrf-1(jam3)* animals that were transformed with either a co-injection marker alone (none) or a co-injection marker along with *rrf-1(+)* expressed under a somatic (*sur-5*), intestine-specific (*sid-2*), or neuron-specific (*rgef-1*) promoter were counted. Red bar, n and asterisks are as in Figure 1C, and ns = not significant.

**Supplementary Figure S3:**
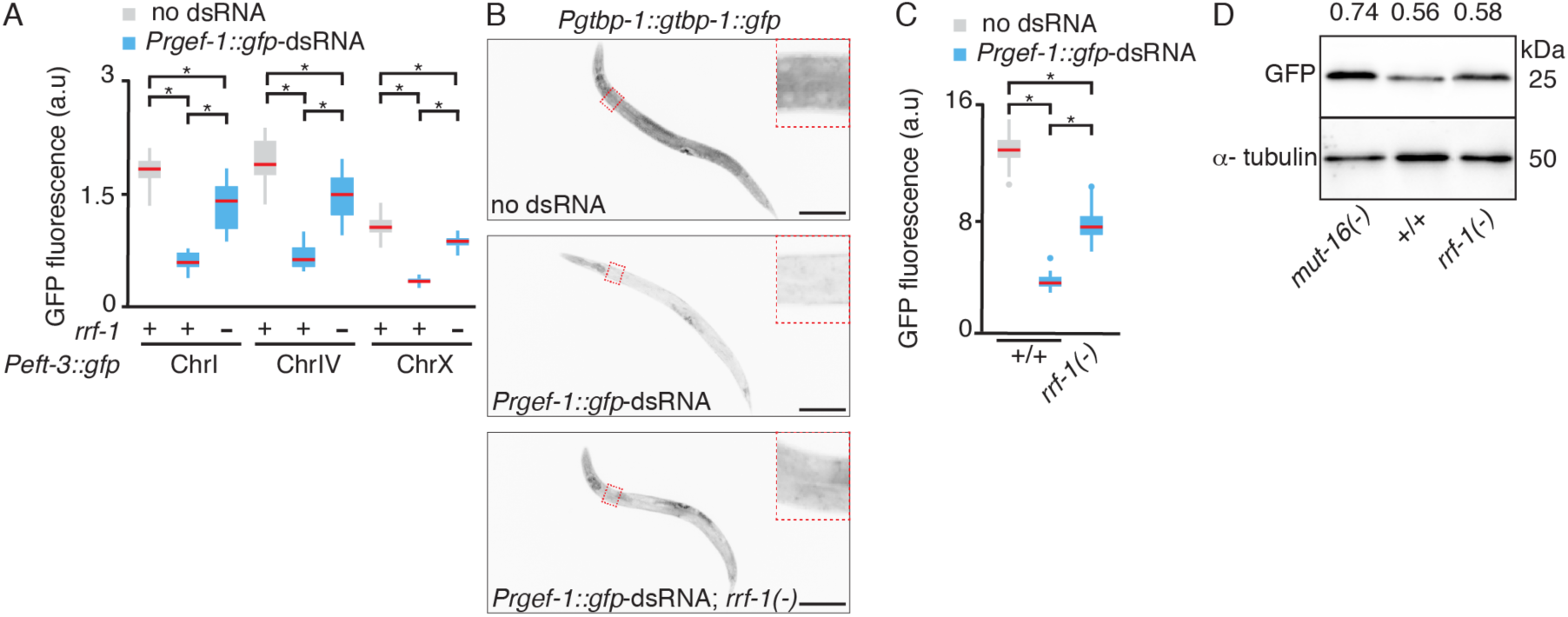
Silencing in the absence of RRF-1 can occur at multiple *gfp* targets expressed under the control of different regulatory elements. (A) Silencing in the absence of RRF-1 does not depend on chromosomal location of target sequences. Effect of *Prgef-1::gfp*-dsRNA and loss of *rrf-1* on GFP fluorescence in animals with *Peft-3::gfp* transgenes located on different chromosomes was quantified as in Figure 2B. Grey boxes, cyan boxes, red bars, n, and asterisks are as in Figure 2B. (B-C) A single-copy gene fusion generated using Cas9-based genome editing can be silenced by neuronal dsRNA in *rrf-1(-)* animals. Representative L4-staged animals that express GFP in all tissues (*Pgtbp-1::gtbp-1::gfp*, *top*) and animals that in addition express *Prgef-1::gfp*-dsRNA in *rrf-1(+)* or *rrf-1(-)* backgrounds (*middle* or *bottom*, respectively) are shown (B). Insets are representative of the region quantified in multiple animals. Quantification of silencing for GFP expressed from *Pgtbp-1::gtbp-1::gfp* is shown (C). Grey boxes, cyan boxes, red bars, n, and asterisks are as in Figure 2B. Scale bar = 50 µm. (D) Silencing in the absence of RRF-1 is associated with a detectable decrease in protein levels. Western blot of GFP protein levels in *Peft-3::gfp* animals expressing *gfp*-dsRNA in an otherwise wild type background (+/+), *mut-16(-)* background (no silencing) or *rrf-1(-)* background (partial silencing). Levels of GFP are normalized to *αα*-tubulin and the median ratios of 3 technical replicates are shown.

**Supplementary Figure S4:**
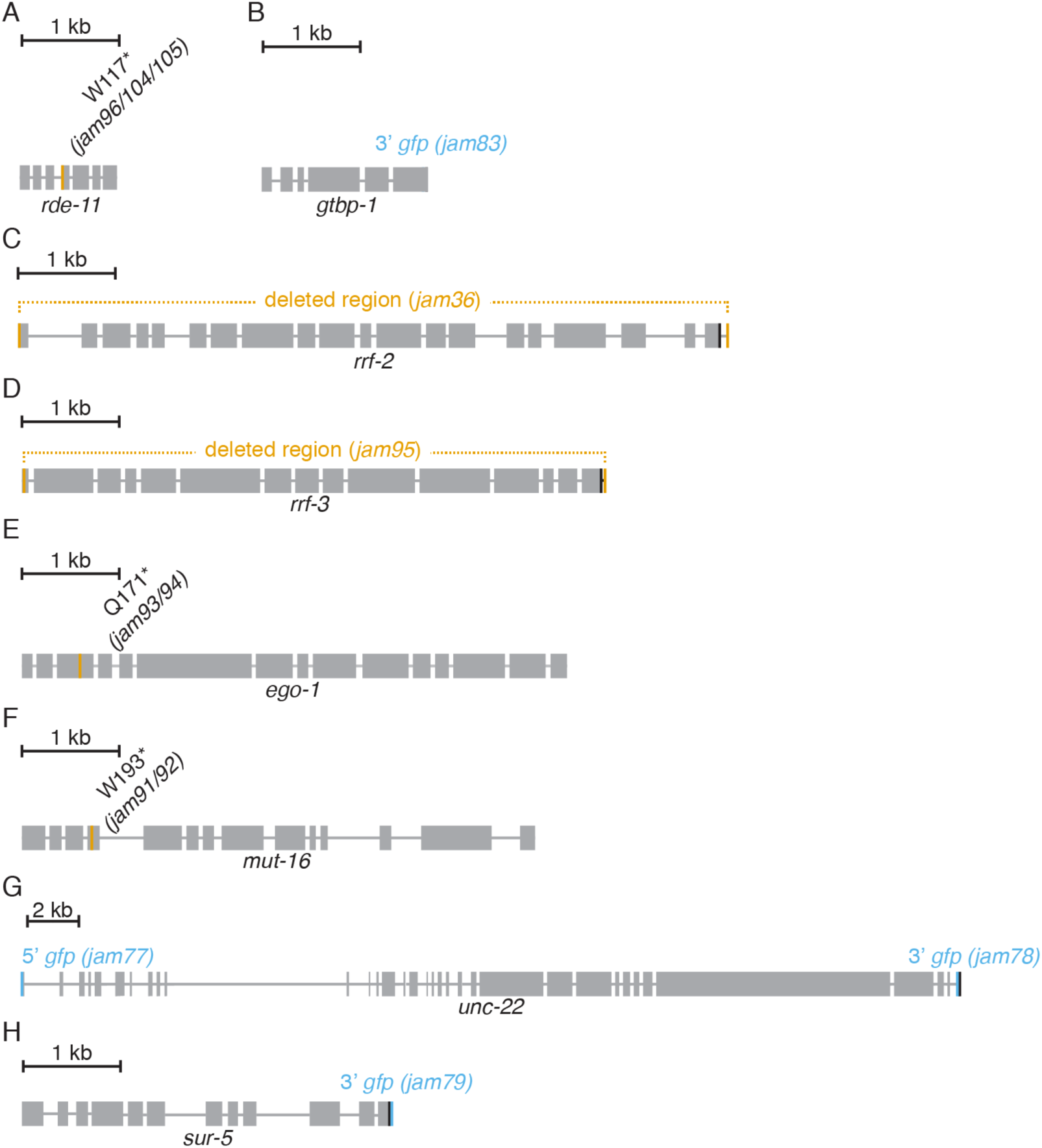
Schematic of genomic changes made using Cas9-based genome editing. Genomic changes introduced into the *rde-11*, *gtbp-1*, *rrf-2*, *rrf-3*, *ego-1*, *mut-16*, *unc-22*, and *sur-5* loci in this study are indicated. The sgRNA target site is indicated (blue for insertions and orange for point mutations and deletions) on the gene structure (exons = grey boxes, introns = grey lines, stop codon = black). Homology-directed repair templates were used to either insert *gfp* sequences (B, G and H), create point mutations (A, E and F) or to delete the region between two target sites (C and D). Asterisks indicate stop codons and scale bars are as indicated.

**Supplementary Figure S5:**
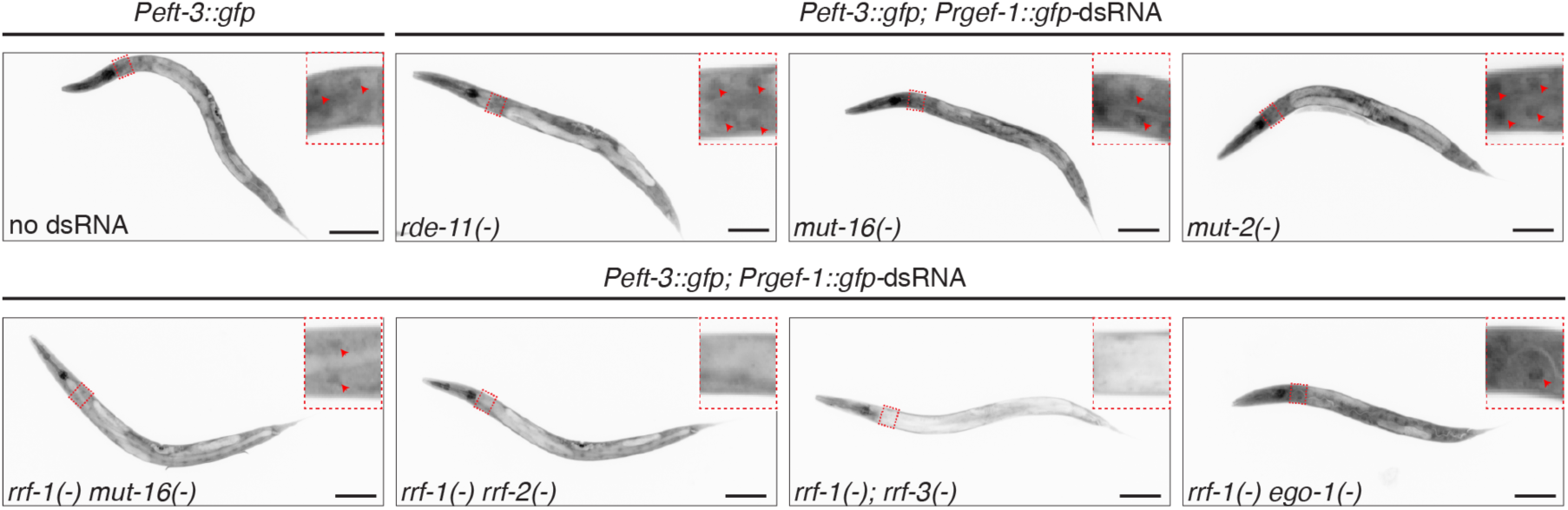
Silencing by neuronal dsRNA in the absence of RRF-1 requires EGO-1 and mutator proteins. Representative images of L4-staged animals expressing no dsRNA or expressing *Prgef-1::gfp*-dsRNA in various single and double mutant backgrounds are shown. Persistent fluorescence in the cytoplasm and nucleus (red arrows) is visible in unsilenced animals (*rde-11(-), mut-16(-), mut-2(-), rrf-1(-) mut-16(-)* and *rrf-1(-) ego-1(-)).* GFP fluorescence in insets are quantified in Figure 2B and 2C.

**Supplementary Figure S6:**
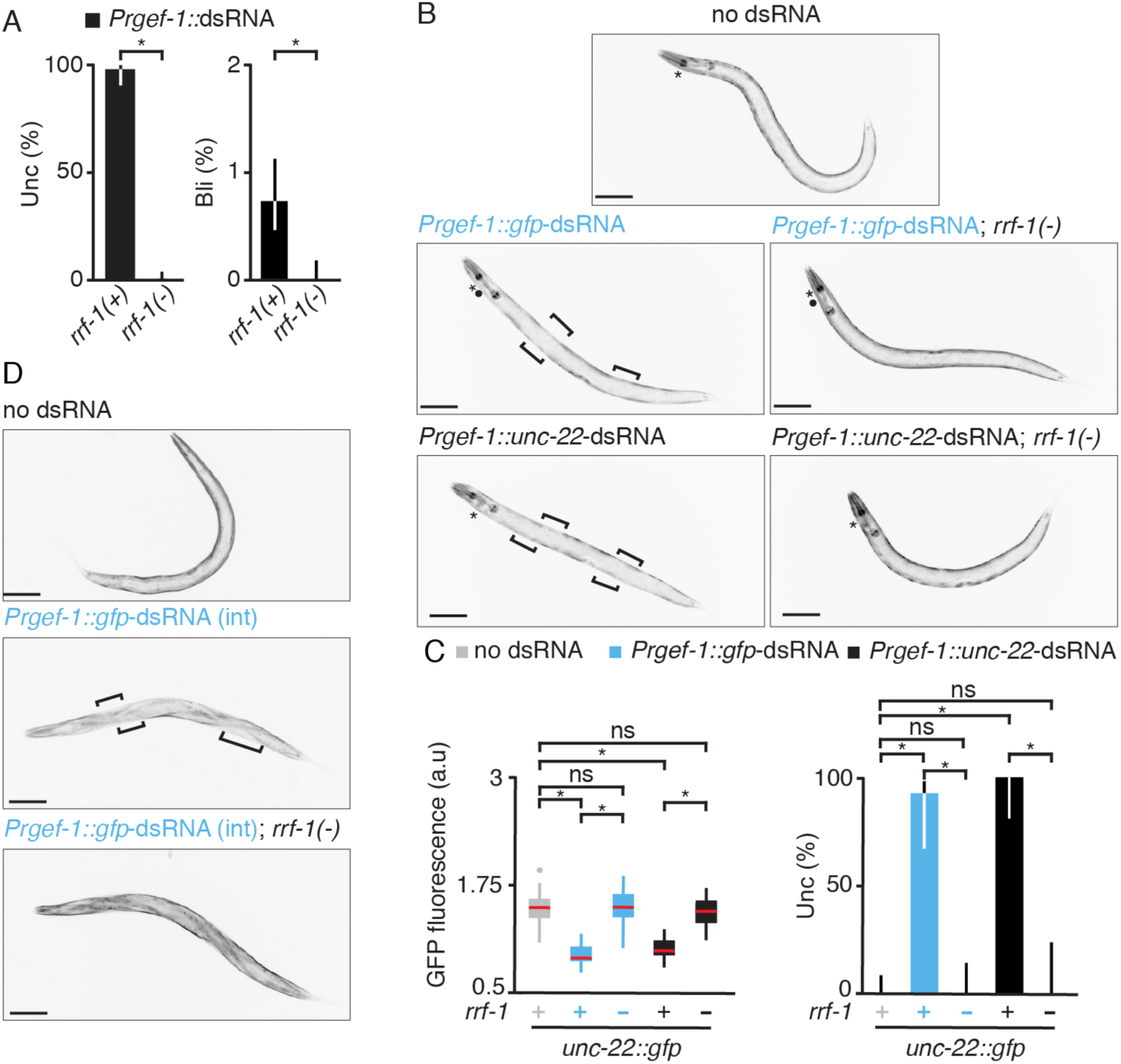
Silencing of a chimeric target by two sources of dsRNA suggests target context and not source of dsRNA dictates RRF-1 requirement. (A) Silencing of *unc-22* or *bli-1* by neuronal dsRNA requires RRF-1. Percentage of animals showing silencing of *unc-22* or *bli-1* by matching dsRNA expressed under a neuronal promoter (*Prgef-1*) was measured in *rrf-1(+)* or *rrf-1(-)* backgrounds. Asterisks indicate p-value < 0.05 (Wilson’s estimates for proportions), n=75 L4-staged animals for *unc-22* and n ≥ 2650 gravid adult animals for *bli-1*. (B-C) Effect of *rrf-1* loss on silencing of a single chimeric target *Punc-22::unc-22::gfp* by either *Prgef-1::gfp*-dsRNA or *Prgef-1::unc-22*-dsRNA was characterized as in Figure 3. Fluorescence in the pharynx is observed in some cases because of expression from *Punc-22::unc-22::gfp* (asterisk, See Materials and Methods) or because of fluorescence from co-injection markers (circle, see Figure legend 3A). (D) An integrated source of neuronal dsRNA that enables silencing in the absence of RRF-1 similarly changes its requirement when used to silence a different target. Representative L4-staged animals that express GFP from *Punc-22::gfp::unc-22* and animals that in addition express *Prgef-1::gfp-*dsRNA (along with a *rol-6* co-injection marker) integrated into a chromosome (int) in *rrf-1(+)* or *rrf-1(-)* backgrounds (*middle* or *bottom*, respectively) are shown. Scale bar = 50 µm.

**Supplementary Figure S7:**
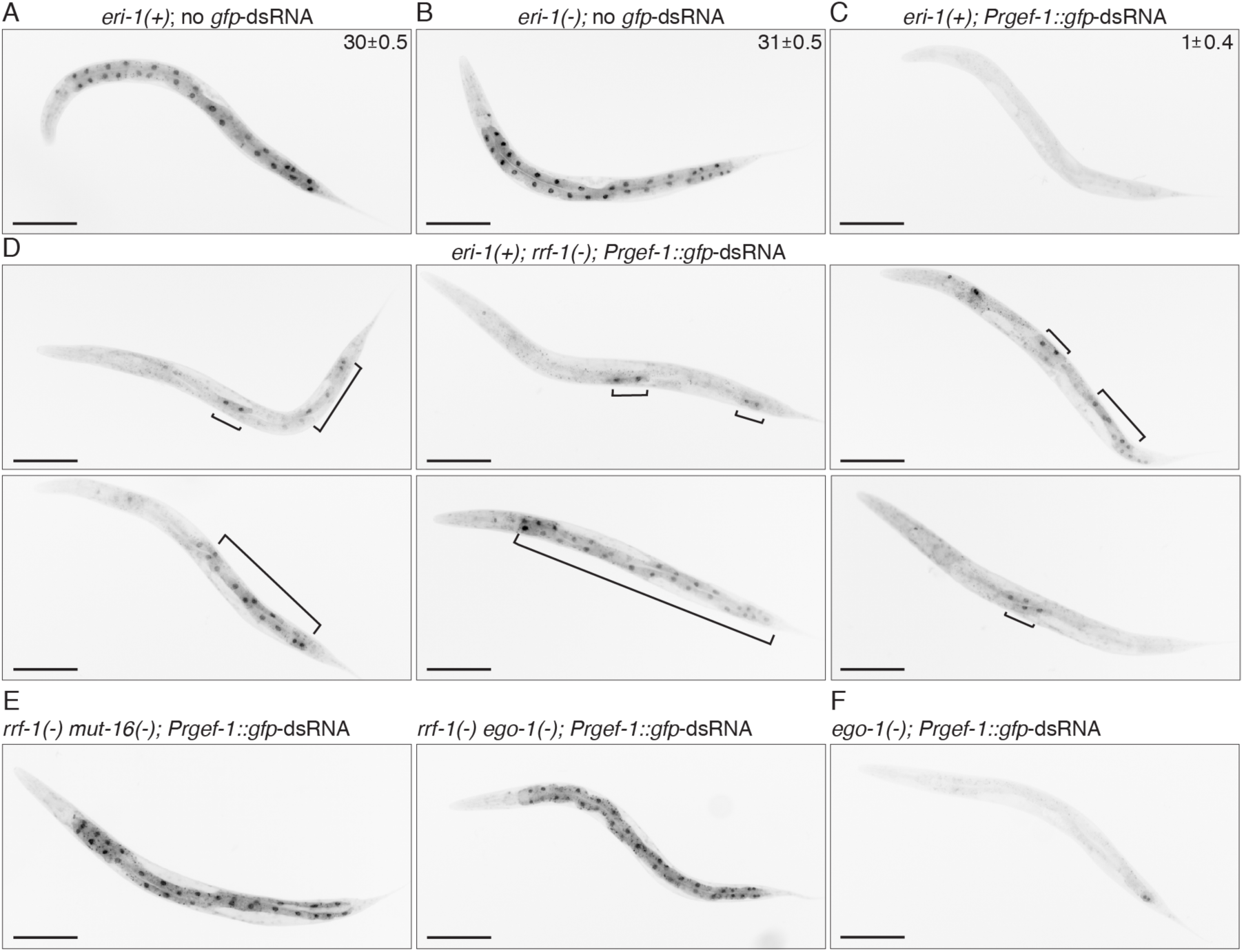
Single-copy *sur-5::gfp* is not subject to self-silencing upon *eri-1* loss but can be robustly silenced by neuronal dsRNA, revealing different patterns of MUT-16- and EGO-1-dependent silencing in an *rrf-1(-)* background. (A-C) Representative *sur-5::gfp* animals in an *eri-1(+)* (i.e., wild-type) background (A) or an *eri-1(-)* background (B) that do not express *gfp*-dsRNA or in an *eri-1(+)* background that express *Prgef-1::gfp-*dsRNA (C) are shown. Average numbers of GFP-positive intestinal nuclei are indicated, errors indicate 95% confidence intervals and n = 25 L4-staged animals. (D) A selection of L4-staged *sur-5::gfp* animals that express *Prgef-1::gfp*-dsRNA in an *eri-1(+); rrf-1(-)* background are shown. Brackets indicate intestinal nuclei that are strongly dependent on RRF-1 for silencing. (E) Silencing in the absence of RRF-1 requires MUT-16 and EGO-1. Representative *sur-5::gfp* animals expressing *Prgef-1::gfp*-dsRNA in *rrf-1(-) mut-16(-)* or *rrf-1(-) ego-1(-)* backgrounds are shown. (F) EGO-1 is required for silencing in intestinal cells only in the absence of RRF-1. A representative *sur-5::gfp;; ego-1(-)* animal expressing *Prgef-1::gfp*-dsRNA is shown. Scale bar = 50 µm in all panels.

**Supplementary Figure S8:**
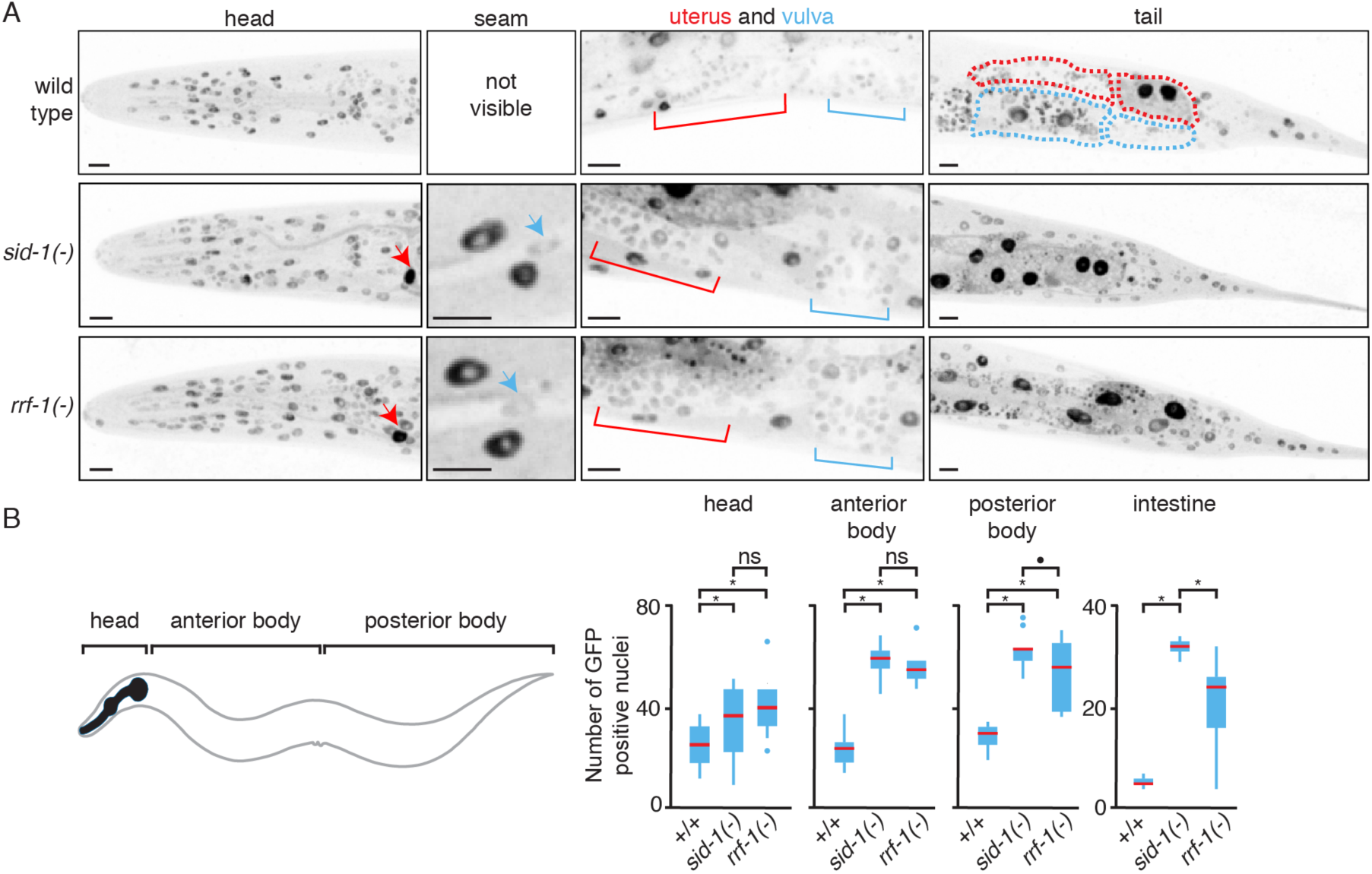
A single-cell resolution view of silencing by neuronal dsRNA reveals differences among cell types and a requirement for RRF-1 in most non-intestinal cells. Silencing of individual nuclei was examined using confocal microscopy of *sur-5::gfp* animals that express *Prgef-1::gfp*-dsRNA in a wild-type, *sid-1(-)*, or *rrf-1(-)* background. (A) Representative images showing maximum intensity projections that highlight extents of silencing in different cell types. The excretory canal cell (red arrow), seam cell (cyan arrow), cells of the developing uterus (red bracket), cells of the developing vulva (cyan bracket), and intestinal cells near the tail of wild-type animals (cyan or red dashed lines for each pair of sister cells) are indicated. Scale bar = 10µm. Seam cells were silenced and not visible in wild-type animals. (B) Silencing by neuronal dsRNA in the absence of RRF-1 is most readily detected in intestinal cells and is highly variable. *Left*, Schematic showing regions where numbers of GFP-positive non-intestinal nuclei were counted (head, anterior to the posterior bulb of the pharynx; anterior body, anterior to the vulva;; posterior body, posterior to the vulva). *Right*, Counts in non-intestinal cells included most body-wall muscle and hypodermal cells throughout the animal but specifically excluded some other cell types. The excretory canal cell (detected in 0/10 *sur-5::gfp*, 10/10 *sid-1(-);; sur-5::gf*p and 10/10 *rrf-1(-);; sur-5::gfp* animals) and intestinal cells were excluded from region-specific analysis based on their large sizes and positions. Cells of the developing uterus and vulva, which remained visible in all tested strains, were excluded based on their small sizes and collective morphology. Seam cells, which show faint expression were excluded because they could not be reliably detected in most Z-slices that were away from the coverslip. Intestinal cells were counted separately. See Supplemental Movie S1 for an example of counting and Materials and Methods for thresholds used. Red bars indicate medians, asterisks and circle indicate p-value < 0.05 and p-value = 0.08, respectively (Student’s t-test), and n = 10 L4-staged animals.

**Supplementary Figure S9:**
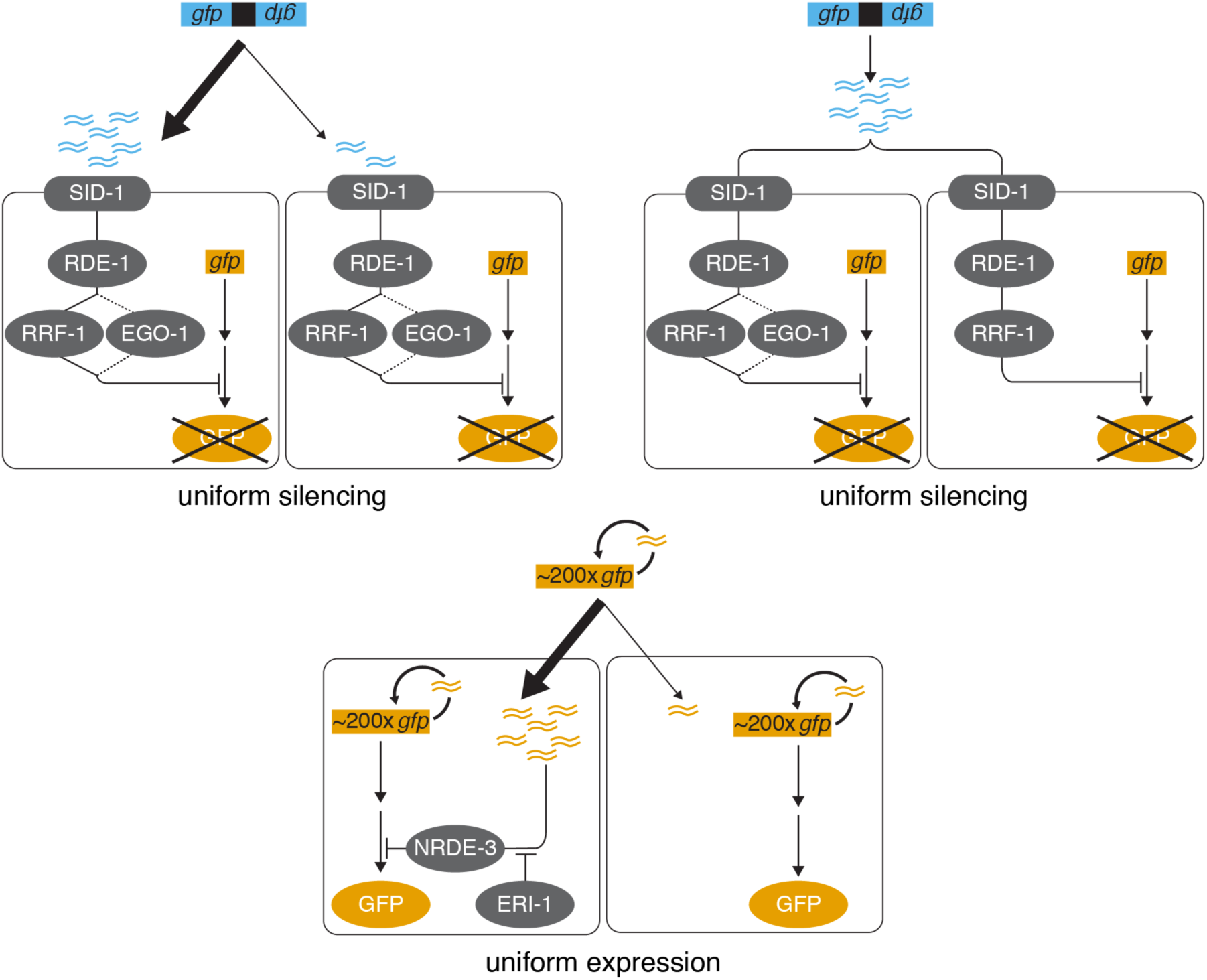
Cells can appear similarly functional despite underlying differences. Two instances where homogeneity of outcome (uniform silencing or uniform gene expression) masks the variation in mechanisms are shown. *Top Left*, Equal amounts of neuronal dsRNA can enter cells and engage either a single RdRP (RRF-1) or two RdRPs (RRF-1 and EGO-1) to eventually cause silencing. In the absence of RRF-1, the unequal and random availability of EGO-1 enables silencing only in some cells. *Top Right*, Equal availability of both RRF-1 and EGO-1 in all intestinal cells could underlie all silencing. Nevertheless, in the absence of RRF-1, each cell amplifies different amounts of small RNAs using EGO-1 because of the unequal and random availability of neuronal dsRNA. Note that silencing enabled by EGO-1 could reflect either direct action in intestinal cells or indirect effects of its expression in the germline. In the presence of RRF-1, this mosaicism is masked and uniform silencing occurs. *Bottom*, Double-stranded RNA derived from a multi-copy *gfp* transgene is segregated asymmetrically during early cell divisions (30). In the absence of ERI-1, the unequal and random availability of this dsRNA prevents expression only in some cells. In the presence of ERI-1, this mosaicism is masked and uniform expression occurs.

Supplementary Movie S1: Z-stack showing counted nuclei (flashes in white) in the anterior body (mostly body-wall muscle and hypodermal cells). See Materials and Methods for the rationale for including/excluding specific nuclei (uterine cells, intestinal cells, etc.).

### SUPPLEMENTARY MATERIALS AND METHODS

#### Transgenesis and genome editing

##### To express *rde-1(+)* under its own promoter (*Prde-1::rde-1(+)*)

The *rde-1* promoter, coding sequence, and 3’ UTR were amplified from N2 gDNA using the primers P17 and P18. AMJ233 animals were transformed with 10 ng/µl of *Prde-1::rde-1(+)* and 40ng/µl of pHC448 in dH_2_O to generate three independent transgenic lines.

##### To express *rde-1(+)* in the neurons (*Prgef-1::rde-1(+)*)

The *rgef-1* promoter (*Prgef-1*) was amplified using the primers P19 and P20, and *rde-1(+)* coding sequence and 3’ UTR was amplified from N2 gDNA using the primers P21 and P22. The two PCR products were used as template and the *Prgef-1::rde-1(+)* fusion product was generated using the primers P23 and P24. AMJ233 animals were transformed with 10 ng/µl of *Prgef-1::rde-1(+)* and 40 ng/µl of pHC448 in dH_2_O to generate three independent transgenic lines.

##### To express *rde-1(+)* in the intestine (*Psid-2::rde-1(+)*)

The *sid-2* promoter (*Psid-2*) was amplified from N2 gDNA using the primers P25 and P26, and *rde-1(+)* coding sequence and 3’ UTR was amplified from N2 gDNA using the primers P27 and P22. The two PCR products were used as template and the *Psid-2::rde-1(+)* fusion product was generated using the primers P28 and P24. AMJ233 animals were transformed with 10 ng/ul of *Psid-2::rde-1(+)* and 40 ng/µl of pHC448 in dH_2_O to generate three independent transgenic lines.

##### To express *rrf-1* in most somatic cells (*Psur-5::rrf-1(+)*)

The precise promoter elements that drive *rrf-1* expression are unclear because *rrf-1* is the downstream gene in an operon that includes another RNA-dependent RNA polymerase gene *ego-1*. Therefore, we used the promoter of a somatically expressed gene *sur-5* to express *rrf-1* in most somatic cells. The *sur-5* promoter was amplified from N2 gDNA using the primers P29 and P30. The *rrf-1* gene and its 3’UTR were amplified together from N2 gDNA using the primers P31 and P32. The two PCR products were used as templates to generate the fusion product using the primers P33 and P34. A 1:4 mixture of *Psur-5::rrf-1(+)* (10 ng/µl) and pHC448 (40 ng/µl) in 10mM Tris HCl (pH 8.5) was injected into AMJ241 to generate AMJ294 and two other independent transgenic lines.

##### To express *rrf-1* in intestinal cells (*Psid-2::rrf-1(+)*)

The *sid-2* promoter was amplified from N2 gDNA using the primers P35 and P36, and the *rrf-1* coding sequence was amplified along with its 3’ UTR using the primers P31 and P32. The two PCR products were used as templates to generate the fusion product using the primers P35 and P34. A 1:4 mixture of *Psid-2::rrf-1(+)* (10 ng/µl) and pHC448 (40 ng/µl) in 10mM Tris HCl (pH 8.5) was injected into AMJ241 to generate AMJ296 and two other independent transgenic lines.

##### To express *rrf-1* in the neurons (*Prgef-1::rrf-1(+)*)

The *rgef-1* promoter was amplified from N2 gDNA using the primers P37 and P38 and the *rrf-1* coding sequence and its 3’UTR were amplified together using the primers P31 and P32. The two PCR products were used as templates to generate the fusion product using the primers P37 and P34. A 1:4 mixture of *Prgef-1::rrf-1(+)* (10 ng/µl) and pHC448 (40 ng/µl) in 10mM Tris HCl (pH 8.5) was injected into AMJ241 to generate AMJ295 and two other independent transgenic lines.

##### To delete *rrf-2* using genome editing

The forward primers P39 and P40 were used to amplify two guide RNAs for the *rrf-2* deletion (Supplementary Figure S4) and the forward primer P41 was used to amplify the guide RNA for the co-conversion marker *dpy-10*. The homology repair templates for *rrf-2* and *dpy-10* were single-stranded DNA oligos (P42 for *rrf-2* and P43 for *dpy-10*). AMJ349 (*Peft-3::gfp;; qtIs49*) animals were injected with 5.1 pmol/µl of *rrf-2* guide RNA1, 6.2 pmol/µl of *rrf-2* guide RNA2, 2.3 pmol/µl of *dpy-10* guide RNA, 9.4 pmol/µl of *rrf-2* homology repair template, 6.1 pmol/µl of *dpy-10* homology repair template and 1.5 pmol/µl of Cas9 protein (PNA Bio Inc.). The deletion was genotyped using 3 primers (P44-P46) and one strain with a homozygous allele was designated as AMJ979.

##### To delete *rrf-3* using genome editing

Two crRNAs were combined with tracrRNA to create an *rrf-3* deletion (Supplementary Figure S4) using a single-stranded DNA homology repair template. AMJ973 animals were injected with 5.0 pmol/µl of *rrf-3* crRNA1(P47) and *rrf-3* guide crRNA2 (P48), 7.2 pmol/µl of tracrRNA (P49), 2.5 pmol/µl of *dpy-10* crRNA (P50), 9.4 pmol/µl of *rrf-3* homology repair template (P51), 9.3 pmol/µl of *dpy-10* homology repair template (P43) and 0.3 pmol/µl of Cas9 protein (IDT). The deletion was genotyped using 3 primers (P52-P54) and one strain with a homozygous allele was designated as AMJ1252.

##### To mutate *mut-16*, *rde-11* and *ego-1* using genome editing

A single crRNA was combined with tracrRNA to target a region of *mut-16*, *rde-11* or *ego-1*. Point mutations in *mut-16* were created in AMJ973 and AMJ977. Point mutations in *rde-11* were created in HC195, HC567 and AMJ301. Point mutations in *ego-1* were created in AMJ973, AMJ976 and AMJ977. Animals were injected with 5 pmol/µl of crRNA (P55 for *mut-16*, P56 for *rde-11* and P57 for *ego-1)*, 4.5-7.2 pmol/µl of tracrRNA, 2.5 pmol/µl of *dpy-10* crRNA (P50), ∼9 pmol/µl of homology repair template (P58 for mut-16, P59 for rde-11 and P60 for ego-1), 9.3 pmol/µl of *dpy-10* homology repair template (P43) and 0.3 pmol/µl of Cas9 protein (IDT). Mutations were genotyped using either the elimination of a restriction site by the point mutation (e.g *rde-11*, genotyped using P05-P06, restriction enzyme: PvuII, strains with a homozygous allele in different backgrounds designated as AMJ1264, AMJ1304 and AMJ1305), or by the addition of silent point mutations that eliminate a nearby restriction site (e.g mut-16, genotyped using P14-P15, restriction enzyme: PstI, strains with a homozygous allele in different backgrounds designated as AMJ974 and AMJ983, and ego-1, genotyped using P61-P62, restriction enzyme: BstBI, strains with a homozygous allele in different backgrounds designated as AMJ1262, AMJ1263 and AMJ1299).

##### To tag *gtbp-1*, *unc-22* and *sur-5* with *gfp* using genome editing

A single guide RNA was selected <6 base pairs away from the insertion site (Supplementary Figure S4), and *gfp* sequences for the homology template was amplified from pTK2 (a derivative of pPD95.75 – Addgene plasmid #1494, a gift from Andrew Fire) using primers with 35-40 base pair overhangs matching either side of the cut site. Guide RNAs were amplified using the forward primers P63 for *gtbp-1::gfp*, P64 for *gfp::unc-22*, P65 for *unc-22::gfp* and P66 for *sur-5::gfp*. Homology templates were amplified using P67 and P68 for *gtbp-1::gfp*, P69 and P70 for *gfp::unc-22*, P71 and P72 for *unc-22::gfp*, and P73 and P74 for *sur-5::gfp*. N2 animals were injected with 9-15 pmol/µl of guide RNA, 0.4 to 0.9 pmol/µl of homology repair template and 1.5 pmol/µl of Cas9 protein (PNA Bio Inc.). Edited F1 or F2 animals were selected by picking animals that showed GFP fluorescence under the Olympus MVX10 fluorescent microscope. The GFP insertion was genotyped using 3 primers (P75 within GFP along with P76 and P77 for *gfp::unc-22*, or P78 and P79 for *unc-22::gfp*, or P80 and P81 for *sur-5::gfp*). Strains with homozygous alleles were designated as AMJ1000 for *gfp::unc-22*, AMJ1001 for *unc-22::gfp* and AMJ975 for *sur-5::gfp*.

**Table S1.**
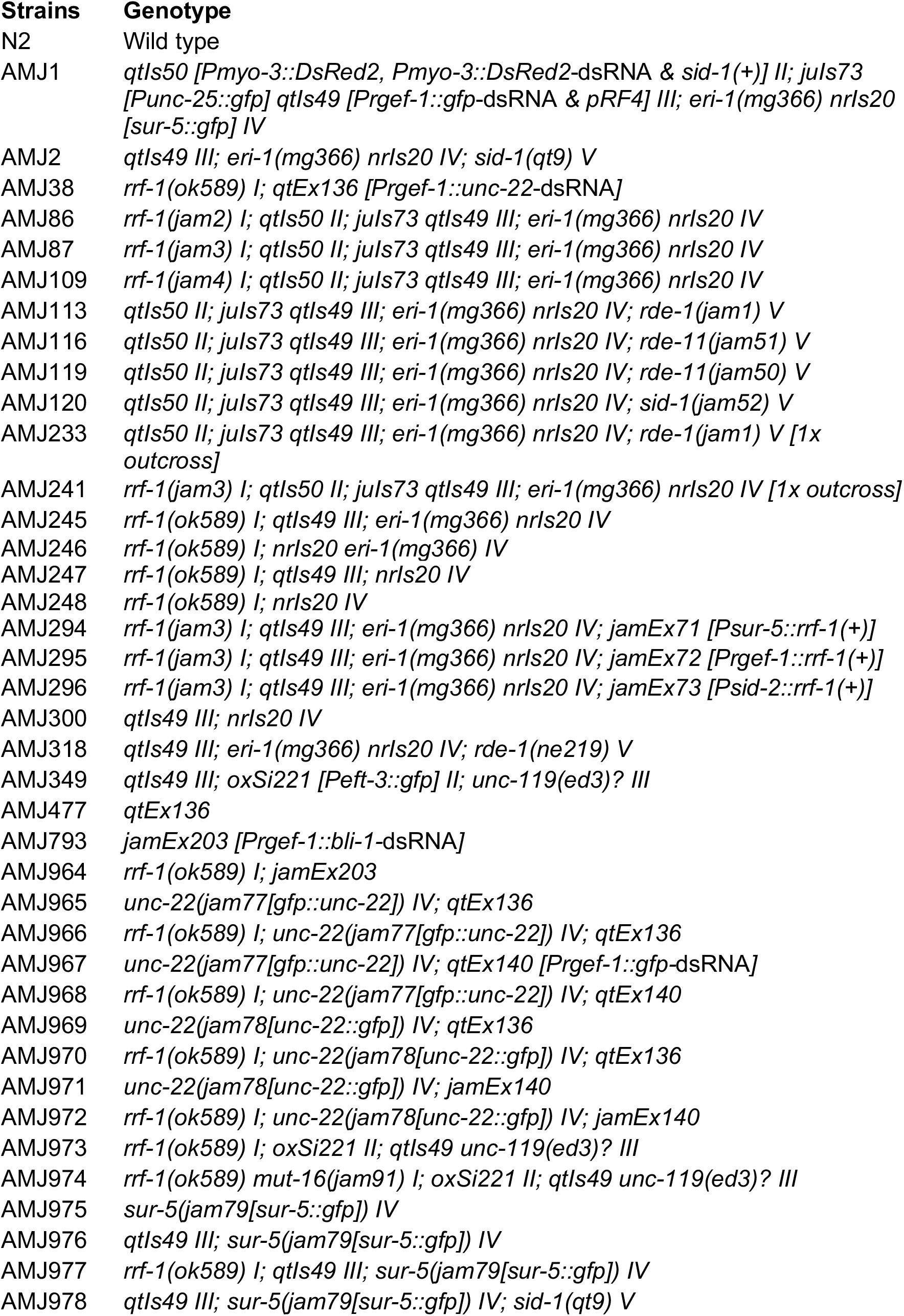
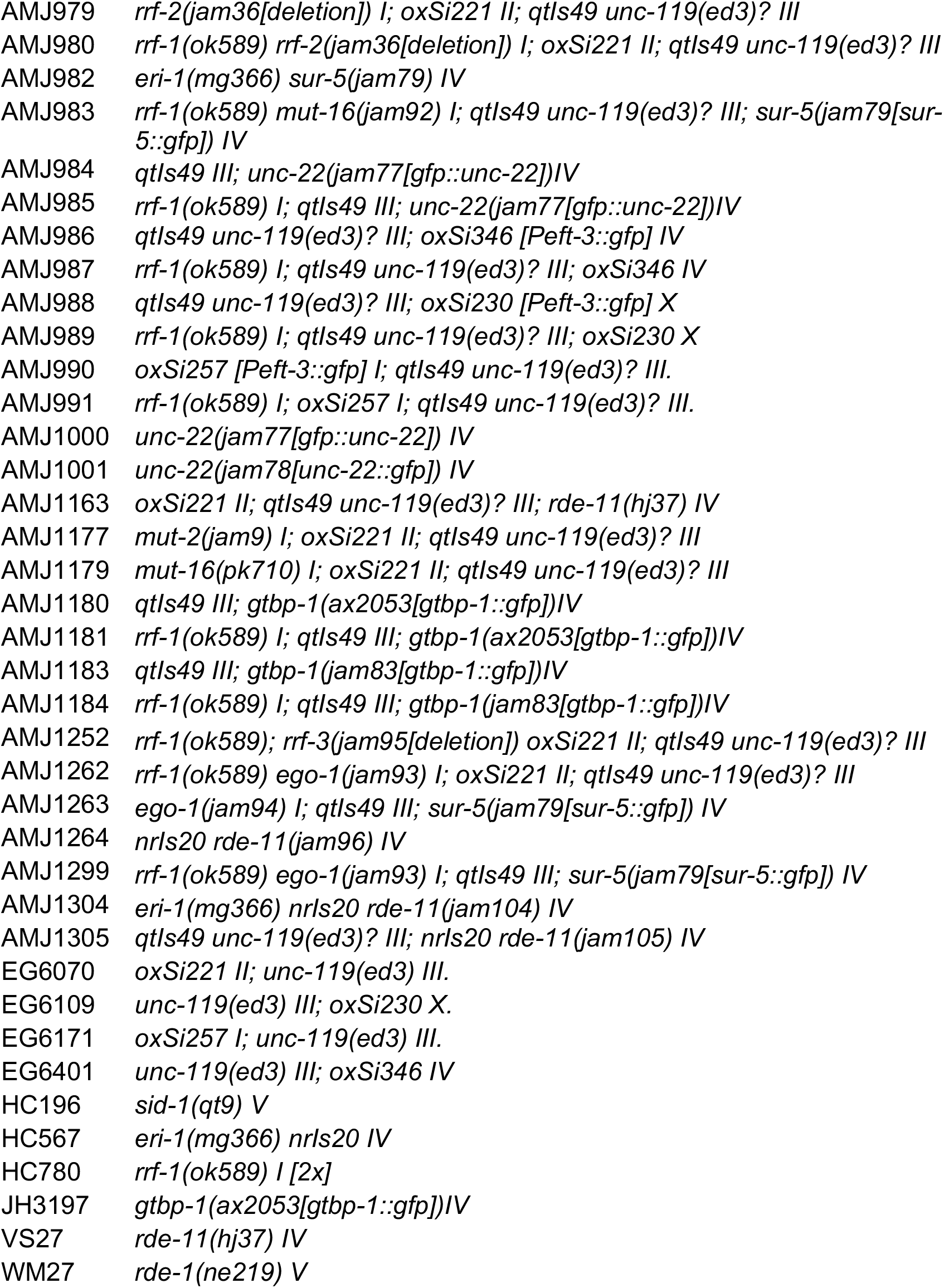
Strains used.

**Table S2.** Sequencing statistics.

| **mutant** | **mapped reads**<br>**(number)** | **mapped**<br>**reads (%)** | **read length**<br>**(bases)** | **genome**<br>**covered (%)** | **depth of**<br>**coverage (fold)** |
| --- | --- | --- | --- | --- | --- |
| *rrf-1(jam2)* | 38,474,029 | 82.8 | 97 | 99.92 | 37.3 |
| *rrf-1(jam3)* | 38,862,516 | 86.8 | 97 | 99.96 | 37.7 |
| *rrf-1(jam4)* | 34,096,083 | 84.0 | 97 | 99.96 | 33.1 |
| *rde-1(jam1)* | 30,401,686 | 81.7 | 97 | 99.96 | 29.5 |
| *rde-11(jam50)* | 42,053,854 | 84.4 | 97 | 99.97 | 40.8 |
| *rde-11(jam51)* | 38,988,725 | 84.0 | 97 | 99.97 | 37.8 |
| *sid-1(jam52)* | 26,823,782 | 73.7 | 97 | 99.95 | 26.0 |

**Table S3.**
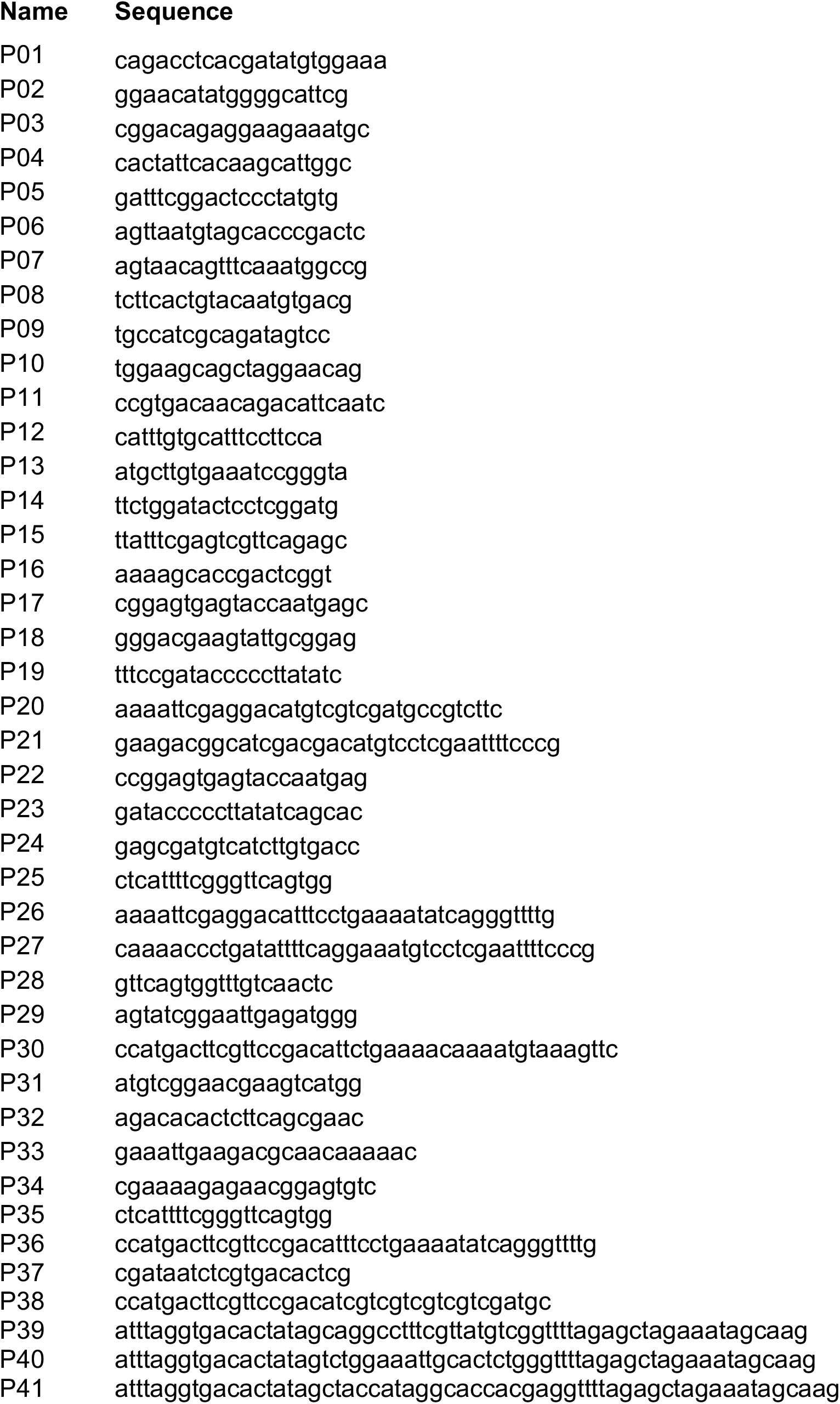
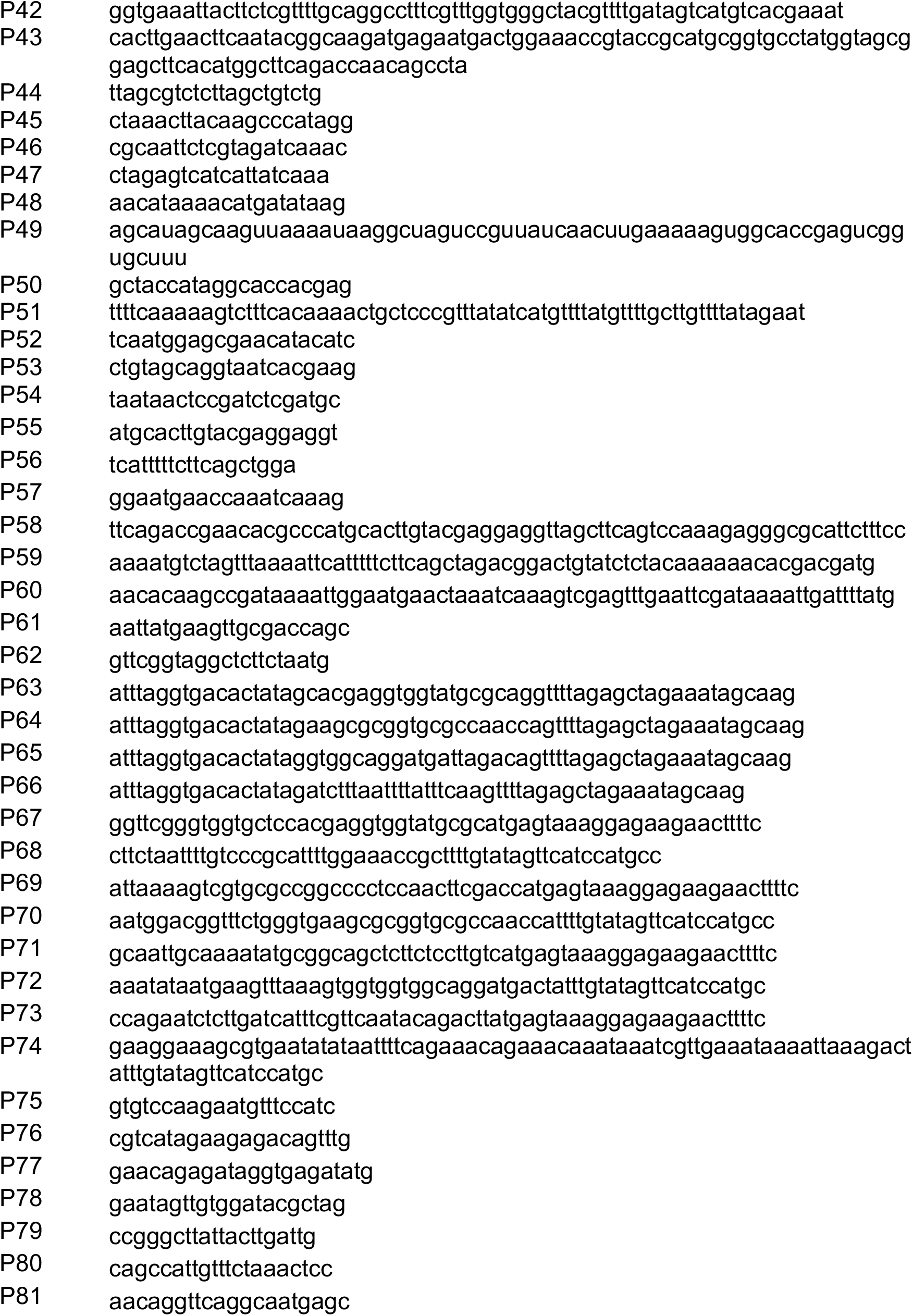
Oligonucleotides used.

## REFERENCES

1. Obbard, D.J., Gordon, K.H., Buck, A.H. and Jiggins, F.M. (2009) The evolution of RNAi as a defence against viruses and transposable elements. Philos Trans R Soc Lond B Biol Sci., 364, 99–115.

2. Fire, A., Xu, S., Montgomery, M.K., Kostas, S.A., Driver, S.E. and Mello, C.C. (1998) Potent and specific genetic interference by double-stranded RNA in *Caenorhabditis elegans*, Nature, 391, 806–11.

3. Perrimon, N., Ni, J.Q. and Perkins, L. (2010) In vivo RNAi: today and tomorrow. Cold Spring Harb Perspect Biol., 2, a003640.

4. Timmons, L. and Fire, A. (1998) Specific interference by ingested dsRNA. Nature, 395, 854.

5. Tabara, H., Grishok, A. and Mello, C.C. (1998) RNAi in *C. elegans*: soaking in the genome sequence. Science, 282, 430–1.

6. Tavernerakis, N., Wang, S.L., Dorovkov, M., Ryazanov, A. and Driscoll, M. (2000) Heritable and inducible genetic interference by double-stranded RNA encoded by transgenes. Nat Genet, 24, 180–3.

7. Winston, W.M., Molodowitch, C. and Hunter, C.P. (2002) Systemic RNAi in *C. elegans* requires the putative transmembrane protein SID-1. Science, 295, 2456–2459.

8. Feinberg, E.H. and Hunter, C.P. (2003) Transport of dsRNA into cells by the transmembrane protein SID-1. Science, 301, 1545–1547.

9. Marré, J., Traver, E.C. and Jose, A.M. (2016) Extracellular RNA is transported from one generation to the next in *Caenorhabditis elegans*. Proc Natl Acad Sci U S A., 113, 12496–12501.

10. Wang, E. and Hunter, C.P. (2017) SID-1 Functions in multiple roles to support parental RNAi in *Caenorhabditis elegans*. Genetics, 207, 547–557.

11. Jose, A.M., Smith, J.J. and Hunter, C.P. (2009) Export of RNA silencing from *C. elegans* tissues does not require the RNA channel SID-1. Proc. Natl. Acad. Sci. U.S.A., 106, 2283–2288.

12. Jose, A.M., Garcia, G.A. and Hunter, C.P. (2011) Two classes of silencing RNAs move between *C. elegans* tissues. Nat. Struct. Mol. Biol., 18, 1184–1188.

13. Raman, P., Zaghab, S.M., Traver, E.C. and Jose, A.M. (2017) The double-stranded RNA binding protein RDE-4 can act cell autonomously during feeding RNAi in *C. elegans*. Nucleic Acids Res., 45, 8463–8473.

14. Devanapally, S., Ravikumar, S. and Jose, A.M. (2015) Double-stranded RNA made in *C. elegans* neurons can enter the germline and cause transgenerational gene silencing. Proc Natl Acad Sci U S A. 112, 2133–8.

15. Tabara, H., Yigit, E., Siomi, H. and Mello, C.C. (2002) The dsRNA binding protein RDE-4 interacts with RDE-1, DCR-1, and a DExH-box helicase to direct RNAi in *C. elegans*. Cell, 109, 861–71.

16. Parker, G.S., Eckert, D.M. and Bass, B.L. (2006) RDE-4 preferentially binds long dsRNA and its dimerization is necessary for cleavage of dsRNA to siRNA. RNA, 12, 807–18.

17. Ketting, R.F., Fischer, S.E., Bernstein, E., Sijen, T., Hannon, G.J. and Plasterk, R.H. (2001) Dicer functions in RNA interference and in synthesis of small RNA involved in developmental timing in *C. elegans*. Genes Dev., 15, 2654–9.

18. Knight, S.W. and Bass, B.L. (2001) A role for the RNase III enzyme DCR-1 in RNA interference and germ line development in *C. elegans*, Science, 293, 2269–71.

19. Tabara, H., Sarikissian, M., Kelly, W.G., Fleenor, J., Grishok, A., Timmons, L., Fire, A. and Mello, C.C. (1999) The *rde-1* gene, RNA interference, and transposon silencing in C. elegans. Cell, 99, 123–32.

20. Pak, J. and Fire, A. (2007) Distinct populations of primary and secondary effectors during RNAi in *C. elegans*. Science, 315, 241–4.

21. Sijen, T., Steiner, F.A., Thijssen, K.L. and Plasterk, R.H. (2007) Secondary siRNAs result from unprimed RNA synthesis and form a distinct class. Science, 315, 244–7.

22. Sijen, T., Fleenor, J., Simmer, F., Thijssen, K.L., Parrish, S., Timmons, L., Plasterk, R.H. and Fire, A. (2001) On the role of RNA amplification in dsRNA-triggered gene silencing. Cell, 107, 465–76.

23. Smardon, A., Spoerke, J.M., Stacey, S.C., Klein, M.E., Mackin, N. and Maine, E.M. (2000) EGO-1 is related to RNA-directed RNA Polymerase and functions in germ-line development and RNA interference in *C. elegans*. Curr Biol., 10, 167–78.

24. Vought, V.E., Ohmachi, M., Lee, M.H. and Maine, E.M. (2005). EGO-1 a putative RNA-directed RNA polymerase, promotes germline proliferation in parallel with GLP-1/notch signaling and regulates the spatial organization of nuclear pore complexes and germline P granules in *Caenorhabditis elegans*. Genetics, 10, 169–78.

25. Guang, S., Bochner, A.F., Pavelec, D.M., Burkhart, K.B., Harding, S., Lachowiec, J. and Kennedy, S. (2008) An Argonaute transports siRNAs from the cytoplasm to the nucleus. Science, 321, 537–541.

26. Ashe, A., Sapetschnig, A., Weick, E.M., Mitchell, J., Bagijn, M.P., Cording, A.C., Doebley, A.L., Goldstein, L.D., Lehrbach, N.J., Le Pen, J., et al. (2012) piRNAs can trigger a multigenerational epigenetic memory in the germline of *C. elegans*. Cell, 150, 88–99.

27. Buckley, B.A., Burkhart, K.B., Gu, S.G., Spracklin, G., Kershner, A., Fritz, H., Kimble, J., Fire, A. and Kennedy, S. (2012) A nuclear Argonaute promotes multigenerational epigenetic inheritance and germline immortality. Nature, 489, 447–51.

28. Shirayama, M., Seth, M., Lee, H.C., Gu, W., Ishidate, T., Conte, D. Jr. and Mello, C.C. (2012) piRNAs initiate an epigenetic memory of nonself RNA in the *C. elegans* germline. Cell, 150, 65–77.

29. Kumsta, C. and Hansen, M. (2012) *C. elegans rrf-1* mutations maintain RNAi efficiency in the soma in addition to the germline. PLoS One, 7, e35428.

30. Le, H.H., Looney, M., Strauss, B., Bloodgood, M. and Jose, A.M. (2016) Tissue homogeneity requires inhibition of unequal gene silencing during development. J Cell Biol., 214, 319–31.

31. Brenner, S. (1974) The genetics of *Caenorhabditis elegans*. Genetics, 77, 71–94.

32. Mello, C.C., Kramer, J.M., Stinchcomb, D. and Ambros, V. (1991) Efficient gene transfer in *C.elegans*: extrachromosomal maintenance and integration of transforming sequences. EMBO J. 10, 3959–70.

33. Dickinson, D.J., Ward, J.D., Reiner, D.J. and Goldstein, B. (2013) Engineering the *Caenorhabditis elegans* genome using Cas9-triggered homologous recombination. Nat Methods, 10, 1028–34.

34. Arribere, J.A., Bell, R.T., Fu, B.X., Artiles, K.L., Hartman, P.S. and Fire, A.Z. (2014) Efficient marker-free recovery of custom genetic modifications with CRISPR/Cas9 in *Caenorhabditis elegans*. Genetics, 198, 837–846.

35. Paix, A., Wang, Y., Smith, H.E., Lee, C.Y., Calidas, D., Lu, T., Smith, J., Schmidt, H., Krause, M.W. and Seydoux, G. (2014) Scalable and versatile genome editing using linear DNAs with microhomology to Cas9 Sites in *Caenorhabditis elegans*. Genetics, 198, 1347–56.

36. Matsunga, Y., Hwang, H., Franke, B., Williams, R., Penley, M., Qudota, H., Yi, H., Morran, L.T., Lu, H., Mayans, O. et al. (2017) Twitchin kinase inhibits muscle activity. Mol Biol Cell, 28, 1591–1600.

37. Goecks, J., Nektrutenko, A., Taylor, J., Galaxy Team. (2010) Galaxy: a comprehensive approach for supporting accessible, reproducible, and transparent computational research in the life sciences. Genome Biol., 11, R86.

38. Blankenberg, D., Von Kuster, G., Coraor, N., Ananda, G., Lazarus, R., Mangan, M., Nekrutenko, A. and Taylor, J. (2010) Galaxy: a web-based genome analysis tool for experimentalists. Curr Proctoc Mol Biol. 19, 1–21.

39. Giardine, B., Riemer, C., Hardison, R.C., Burhans, R., Elnitski, L., Shah, P., Zhang, Y., Blankenberg, D., Albert, I., Taylor, J. et al. (2005) Galaxy: a platform for interactive large-scale genome analysis. Genome Res., 15, 1451–5.

40. Minevich, G., Park, D.S., Blankenberg, D, Poole, R.J. and Hobert, O. (2013) CloudMap: a cloud-based pipeline for analysis of mutant genome sequences. Genetics., 192, 1249–69.

41. Miller, R.G., Jr. (1981) Simultaneous statistical inference. McGraw-Hill, New York, NY, USA.

42. Lee, C., Sorensen, E.B., Lynch, T.R. and Kimble, J. (2016b). *C. elegans* GLP-1/Notch activates transcription in a probability gradient across the germline stem cell pool, eLife, 5,e18370.

43. Lee, C., Seidel, H.S., Lynch, T.R., Sorensen, E.B., Crittenden, S.L. and Kimble, J. (2017). Single-molecule RNA Fluorescence *in situ* Hybridization (smFISH) in *Caenorhabditis elegans*. Bio-protocol, 7, e2357.

44. Schindelin, J., Arganda-Carreras, I., Frise, E., Kaynig, V., Longair, M., Pietzsch, T., Preibisch, S., Rueden, C., Saalfeld, S., Schmid, B. et al. (2012) Fiji: an open-source platform for biological-image analysis. Nat Methods, 9, 676–82.

45. Preibisch, S., Saalfeld, S. and Tomancak, P.. (2009) Globally optimal stitching of tiled 3D microscopic image acquisitions. Bioinformatics, 25,1463–5.

46. Bolte, S. and Cordelières, F.P. (2006) A guided tour into subcellular colocalization analysis in light microscopy. J Microsc, 224, 213–32.

47. Sulston, J.E. and Horvitz, H.R. (1977) Post-embryonic cell lineage of the nematode, *Caenorhabditis elegans*. Dev. Biol., 56, 110–156.

48. Sulston, J.E., Schierenberg, E., White, J.G, and Thompson, J.N. (1983) The embryonic cell lineage of the nematode *Caenorhabditis elegans*. Dev. Biol., 100, 64–119.

49. Leung, B., Hermann, G.J. and Priess, J.R. (1999) Organogenesis of the *Caenorhabditis elegans* intestine. Dev. Biol., 216, 114–134.

50. Hermann, G.J., Leung, B. and Priess, J.R. (2000) Left-right asymmetry in C. elegans intestine organogenesis involves a LIN-12/Notch signaling pathway. Development, 127, 3429–3440.

51. Mendenhall, A. R., Tedesco, P.M., Sands, B., Johnson, T.E. and Brent, R. (2015) Single cell quantification of reporter gene expression in live adult *Caenorhabditis elegans* reveals reproducible cell-specific expression patterns and underlying biological variation. PLoS One, 10, e0124289.

52. Asan, A., Raiders, S. A. and Priess, J. R. (2016) Morphogenesis of the *C. elegans* Intestine Involves Axon Guidance Genes. PLoS Genet. 12, e1005950.

53. Newcombe, R.G. (1998) Two-sided confidence intervals for the single proportion: comparison of seven methods. Statist. Med., 17, 857–872.

54. Grishok, A.(2013) Biology and mechanisms of short RNAs in *Caenorhabditis elegans*. *Adv*. Genetics, 83, 1–69.

55. Hellwig, S. and Bass, B.L. (2008) A starvation-induced noncoding RNA modulates expression of Dicer-regulated genes. Proc. Natl. Acad. Sci. U.S.A., 105, 12897–12902.

56. Zhang, C., Montgomery, T.A., Fischer, S.E., Garcia, S.M., Riedel, C.G., Fahlgren, N., Sullivan, C.M., Carrington, J.C. and Ruvkun, G. (2012) The *Caenorhabditis elegans* RDE-10/RDE-11 complex regulates RNAi by promoting secondary siRNA amplification. Curr Biol., 22, 881–90.

57. Yang, H., Zhang, Y., Vallandingham, J., Li, H., Floren, L. and Mak, H.Y. (2012) The RDE-10/RDE-11 complex triggers RNAi-induced mRNA degradation by association with target mRNA in *C. elegans*. Genes Dev., 26, 846–56.

58. Vastenhouw, N.L., Fischer, S.E., Robert, V.J., Thijssen, K.L., Fraser, A.G., Kamath, R.S., Ahringer, J and Plasterk, R.H. (2003) A genome-wide screen identifies 27 genes involved in transposon silencing in *C. elegans*. Curr Biol., 13, 1311–6.

59. Chen, C.C., Simard, M.J., Tabara, H., Brownell, D.R., McCollough, J.A. and Mello, C.C. (2005) A member of the polymerase beta nucleotidyltransferase superfamily is required for RNA interference in *C. elegans*. Curr Biol., 15, 378–83.

60. Phillips, C.M., Montgomery, T.A., Breen, P.C. and Ruvkun, G. (2012) MUT-16 promotes formation of perinuclear mutator foci required for RNA silencing in the *C. elegans* germline. Genes Dev., 26, 1433–44.

61. Uebel, C.J., Anderson, D.C., Mandarino, L.M., Manage, K.I., Aynaszyan, S. and Philips, C.M. (2018) Distinct regions of the intrinsically disordered protein MUT-16 mediate assembly of a small RNA amplification complex and promote phase separation of Mutator foci. PLoS Genet, 14, e1007542.

62. Sarkies, P., Selkirk, M.E., Jones, J.T., Blok, V., Boothby, T., Goldstein, B., Hanelt, B., Ardilia-Garcia, A., Fast, N.M., Schiffer, P.M. et al. (2015) Ancient and novel small RNA pathways compensate for the loss of piRNAs in multiple independent nematode lineages. PLoS Biol., 13, e1002061.

63. Blumenfeld, A. and Jose, A.M. (2016) Reproducible features of small RNAs in *C. elegans* reveal NU RNAs and provide insights into 22G RNAs and 26G RNAs. RNA, 22,184–192.

64. Gent, J.I., Lamm, A.T., Pavelec, D.M., Maniar, J.M., Parameswaran, P., Tao, L., Kennedy, S. and Fire, A.Z. (2010) Distinct phases of siRNA synthesis in an endogenous RNAi pathway in *C. elegans* soma. Mol Cell., 37, 679–89.

65. Simmer, F., Tijsterman, M., Parrish, S., Koushika, S.P., Nonet, M.L., Fire, A., Ahringer, J. and Plasterk, R.H. (2002) Loss of the putative RNA-directed RNA polymerase RRF-3 makes *C. elegans* hypersensitive to RNAi. Curr. Biol., 12, 1317–1319.

66. Maniar, J.M. and Fire, A.Z. (2011) EGO-1, a *C. elegans* RdRP, modulates gene expression via production of mRNA-templated short antisense RNAs. Curr Biol., 21, 449–59.

67. Pak, J., Maniar, J.M, Mello, C.C. and Fire, A. (2012) Protection from feed-forward amplification in an amplified RNAi mechanism. Cell, 151, 885–99.

68. Calixto, A., Chelur, D., Topalidou, I., Chen, X. and Chalfie, M. (2010) Enhanced neuronal RNAi in *C. elegans* using SID-1. Nat. Methods, 7, 554–559.

69. Lee, R.C., Hammel, C.M. and Ambros, V. (2006) Interacting endogenous and exogenous RNAi pathways in *Caenorhabditis elegans*. RNA, 12, 589–97.

70. Whipple, J.M., Youssef, O.A., Aruscavage, P.J., Nix, D.A., Hong, C., Johnson, W.E. and Bass, B.L. (2015) Genome wide profiling of the *C. elegans* dsRNAome. RNA, 21, 786–800,

71. Zhang, H. and Fire, A.Z. (2010) Cell autonomous specification of temporal identity by *Caenorhabditis elegans microRNA lin-4*. Dev Biol, 344, 603–10.

72. Gu, W., Shirayama, M., Conte, D. Jr., Vasale, J., Batista, P.J., Claycomb, J.M., Moresco, J.J., Youngman, E.M., Keys, J., Stoltz, M.J., et al. (2009) Distinct agonuate-mediated 22G-RNA pathways direct genome surveillance in the *C. elegans* germline. Mol Cell, 36, 231–44.

73. Waldron, F.M., Stone, G.N. and Obbard, D.J. (2018) Metagenomic sequencing suggests a diversity of RNA interference-like responses to viruses across multicellular eukaryotes. PLoS Genet, 14, e1007533.

74. Félix, M.A., Ashe, A., Piffaretti, J., Wu, G., Nuez, I., Bélicard, T., Jiang, Y., Zhao, G., Franz, C.J., Goldstein, L.D. et al. (2011) Natural and experimental infection of *Caenorhabditis* nematodes by novel viruses related to nodaviruses. PLoS Biol, 9, e1000586.

75. Franz, C.J., Renshaw, H., Frezal, L., Jiang, Y., Félix, M.A. and Wang, D. (2014) Orsay, Santeuil and Le Blanc viruses primarily infect intestinal cells in *Caenorhabditis* nematodes. Virology, 5, 255–64.

76. Buganim, Y., Faddah, D.A., Cheng, A.W., Itskovich, E., Markoulaki, S., Ganz, K., Klemm, S.L., Van Ourdenaarden, A. and Jaenisch, R. (2012) Single-cell expression analyses during cellular reprogramming reveal an early stochastic and late hierarchic phase. Cell, 150, 1209–22.

77. Liberali, P., Snijder, B. and Pelkmans, L. (2014) A hierarchical map of regulatory genetic interactions in membrane trafficking. Cell, 157, 1473–87.

78. Gut, G., Hermann, M.D. and Pelkmans, L. (2018) Multiplexed protein maps link subcellular organization to cellular states. Science, 361, eaar7042.

